# Integrating structural modeling and divergence dating of RNA-dependent RNA polymerases to resolve the evolutionary history of plant and fungal viruses: from sobemoviruses to sobelivirads

**DOI:** 10.1101/2024.11.13.623411

**Authors:** Jérôme Gracy, Mahan Ghafari, Gilles Labesse, Denis Fargette, Eugénie Hébrard

## Abstract

RNA-dependent RNA polymerases (RdRps) are crucial for RNA virus replication and serve as key marker genes for defining deep taxonomic ranks and for understanding viral evolutionary history. Despite their conserved functions and motifs, the high genetic diversity of RdRps complicates precise sequence comparisons across viral families, hindering accurate taxonomic classification of new species. Three-dimensional (3D) RdRp structures can help overcome these challenges through structure-guided alignments. However, such data are rare for myco- and phytoviruses, limiting investigation of their ecological and evolutionary links. In this study, we focused on the highly divergent order *Sobelivirales*, which includes sobemoviruses infecting plants — a taxon known for its ancient origin — and barnaviruses infecting fungi, which remain relatively unknown. Using deep-learning structural modeling, we generated reliable 3D models for 44 sobemoviral and sobelivirad species. Structure-guided alignments, together with new barnaviral and relevant outgroup sequences, enabled robust phylogenetic reconstruction, allowing us to propose revisions of existing viral families and suggest new evolutionary scenarios. Divergence dates were estimated for the first time at this taxonomical rank using the Prisoner of War model, which revealed a divergence of plant and fungal sobelivirads 27.0 ± 10 million years ago — much more recent than the separation of their respective hosts. This result suggests that cross-kingdom host shifts have contributed more likely to the evolutionary history of *Sobelivirales* than strict virus-host codivergence. Based on an extended dataset of 127 species, structure and sequence conservation analyses identified molecular signatures of sobeliviral families. These conserved and extented RdRp motifs will facilitate future taxonomic assignments and the development of diagnostic tools. Our interdisciplinary approach, integrating structure modeling and divergence dating, offers new insights into the evolutionary divergence of plant and fungal viruses, with potential applications to other viral orders and families.

**Author summary:** RNA-dependent RNA polymerases (RdRps) are essential for RNA virus replication and serve as important markers for classifying viruses and understanding their evolution. However, with metagenomic studies that rapidly expand the known diversity of RNA viruses, it is increasingly difficult to compare highly divergent RdRps and accurately classify new species. When available, 3D structures of RdRps can help overcome this challenge through structure-guided alignments.

Here, we focused on the highly divergent order *Sobelivirales*, which groups phytoviruses and mycoviruses. Deep-learning models were used to generate reliable 3D structures for 44 representative viral species. Structure-guided alignments combined with the identification of new barnaviruses and outgroups allowed us to build a more accurate viral phylogeny. Based on this finding, we proposed updates to existing viral families and genera within the order *Sobelivirales*. We estimated divergence dates using a model that previously uncovered the ancient origins of sobemovirus. Notably, we provided the first estimate of when these plant and fungal viruses diverged ∼27.0 ± 10 million years ago, suggesting cross-kingdom host shifts rather than strict virus-host codivergence. We also identified molecular signatures that are useful for future virus classification and diagnosis, with potential applications to other viral groups.

## Introduction

Recent advances in viral metagenomics have revealed how extensive and unexplored viral diversity is, and have contributed significantly to a better understanding of viral evolution. Major adaptations of viral taxonomic classification have been made to incorporate numerous new lineages and their evolutionary links [1,2]. This includes the creation of novel upper ranks and subdivisions such as classes and phyla [3]. The identification of viral hallmark genes (VHG) facilitated the assignment of viral taxa into these novel groups [4,5]. Polymerases especially RNA-dependent RNA polymerases (RdRp) represent one of the major groups of VHG due to their crucial function for realm *Riboviria* (RNA viruses) [6]. RdRp comparative analyses revealed unexpected relationships between some viral groups, prompting taxonomic reassessments and modifications. For example, the family *Luteoviridae* had initially grouped the genera *Luteovirus* and *Polerovirus*, primarly due to their similar circulative transmission mode mediated by aphids and the related readthrough domain of their capsid protein (CP). However, based on the low sequence similarity of their RdRps, the family *Luteoviridae* was abolished by the International Committee on Taxonomy of Viruses (ICTV) and the two genera were subsequently reassigned to the families *Tombusviridae* and *Solemoviridae*, respectively [7].

Viral RdRps are monomeric proteins responsible for synthesizing complementary RNA strands from a template RNA. They exhibit an overall 3D structure often likened to a cupped human right hand in which the fingers and thumb domains interact to form a characteristic central tunnel (for review [8–11]). The palm domain is formed by a β-sheet with three to six β-strands above two α-helices, the fingers domain is a mixed α/β structure and the thumb domain is highly variable with a predominantly helical structure. The fingers and thumb domains contribute to the RdRp overall stability and to proper positioning of the RNA template and nascent RNA strand. The palm domain is essential for the accurate and efficient RNA chain elongation. It allows (i) the positioning of the primer 3’OH, or the nucleotidylated viral protein genome-linked (VPg) depending on the viral taxon, (ii) the binding of incoming nucleotides and (iii) their incorporation to the RNA chain using two metal ions as cofactors. Comparisons of RdRp sequences of diverse viral groups has identified short linear motifs of highly conserved amino acids [12–14]. Five motifs (A-E) were reported in the palm domain, two motifs (F and G) in the fingers domain and one putative and less conserved motif (H) in the thumb domain. High-throughput taxonomic assignments were often based on the ABC motifs only [15–17]. Analyzing the 3D structures of RdRps later identified larger protein segments referred to as homomorphs, determined by structural similarities rather than sequence composition [18]. Enlarging the dataset of available 3D structures and releasing sequence conservation constraints allowed the identification of a common structural core and ancient phylogenetic relationships between distants taxa [19]. However, for some viral phyla such as those of plant and fungus viruses, the absence of 3D structures poses problems for comparing RdRps and revealing their evolutionary history. To address this challenge, multiple protein alignments were restricted to the central RdRp palm domain, using classical algorithms combined with manual refinements based on available 3D structures from taxonomically distant related viruses, such as animal or human viruses. These approaches led to the creation of the kingdom *Orthornavirae* and its associated phyla [4,6]. However, these approaches lack the precision needed to analyze in detail phylogenetic relationships between lower taxonomic ranks such as classes, orders and families.

Although the monophyly of viruses in the kingdom *Orthornavirae* has been established based on comparison of their RdRps, reconstruction of their evolutionary history remains difficult due to their strong, probably ancient, evolutionary divergence and the central role of horizontal transfer of other viral genes in plant viruses [20]. In the picorna-like supergroup, now referred as the phylum *Pisuviricota*, viral species share common structural and functional features in RdRp. On the other hand, they also exhibit contrasting biological characteristics in terms of host range (animals, insects, plants, fungi, protists), as well as variations in genomic molecule (positive single-stranded RNA or double-stranded RNA) and size (ranging from the largest to the smallest RNA genomes) (S1 Table). Phylogenetic analyses of viral taxa, such as the phylum *Pisuviricota* or its main class *Pisoniviricetes,* can reveal evolutionary relationships that help formulate hypotheses about the origins of shared or distinct traits among these viruses.

Viruses of fungi also named mycoviruses have long been understudied, but advances of high-throughput sequencing technologies have revealed the large and unexpected diversity of fungal viruses [21,22]. Moreover, recent discoveries, such as cross-kingdom transmission events, highlight ecological and evolutionary links with plant viruses [20,22–24]. In this study, we compared the RdRps inside the order *Sobelivirales*, as markers of their common history to gain insights into host range evolution. This order is one of the six orders of positive-sense single-stranded RNA (+ssRNA) mycoviruses. It comprises families *Solemoviridae, Barnaviridae* and *Alvernaviridae*, which contains plant, fungus and micro-algae viruses, respectively. Solemovirids (members of the family *Solemoviridae*) includes four plant viral genera (*Sobemovirus*, *Polerovirus*, *Enamovirus* and *Polemovirus*), while barnavirids and alvernavirids consist of single genera, *Barnavirus* and *Dinornavirus*, each composed of only one approved species infecting fungus or dinoflagellates, respectively (Table 1). Apart from the alvernavirids which are not yet well known, these families share the same characteristic genomic arrangement of their main polyprotein such as chymotrypsine-like protease / VPg / RdRp [25,26], but their sequence conservation is restricted to very short linear motifs. Similarities between RdRp active sites of sobemovirus and polero-/enamovirus were noted as early as 1991 [12] — leading finally to their taxonomic grouping in 2021 [27]. Although RdRp homologies between sobemoviruses and barnaviruses were proposed in 2008 [10], they have not yet resulted in taxonomic revisions. Furthermore, a newly proposed genus of fungal virus, *Hubsclerovirus*, has recently been assigned to the phytoviral family *Solemoviridae* instead of the mycoviral family *Barnaviridae* [28]. Thus, comparing more precisely the RdRps from these viral families would provide valuable insights into the complex history of plant and fungus viruses and would contribute to revise their taxonomy.

**Table 1:**
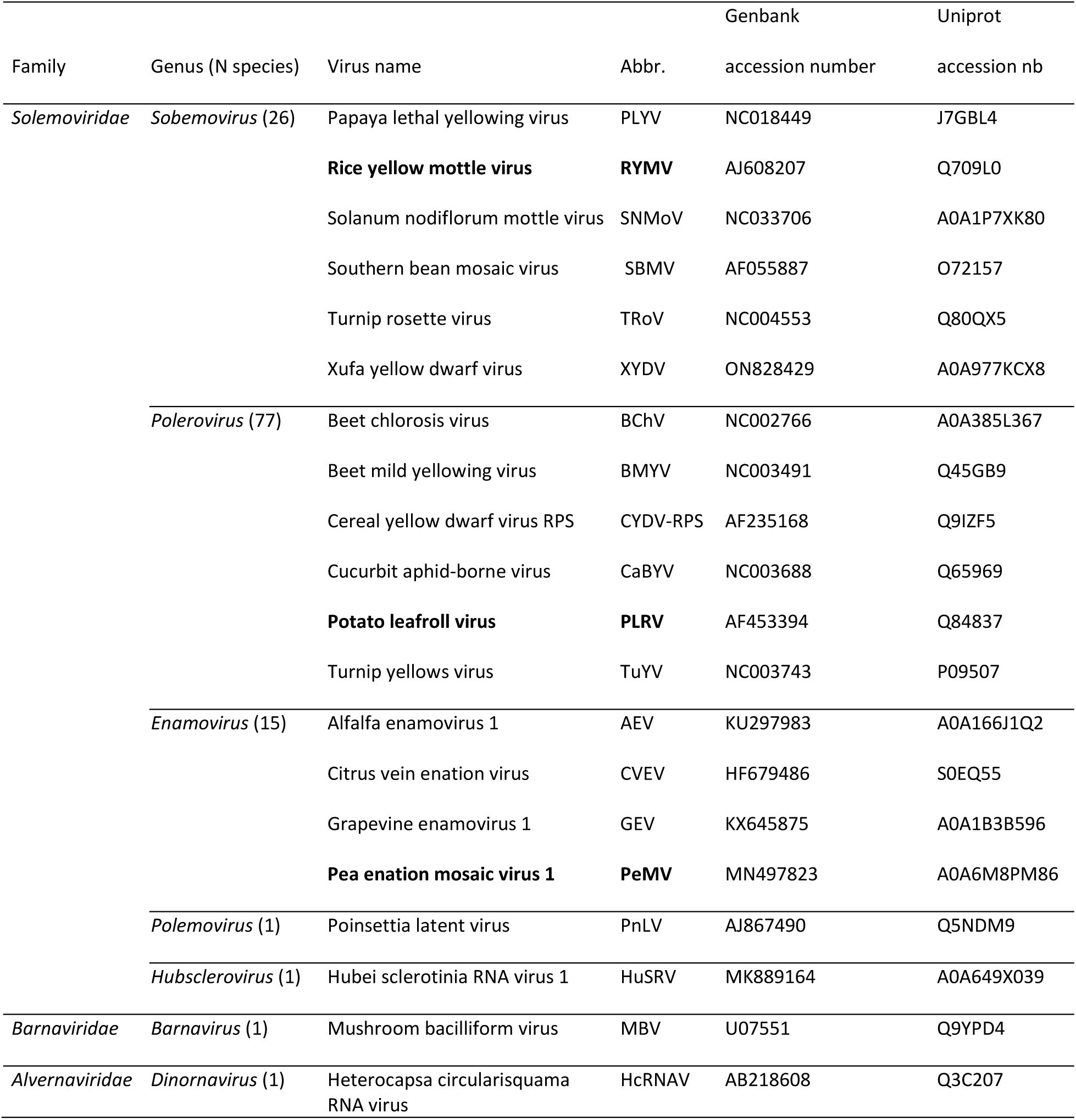
Composition of the order *Sobelivirales* and species analyzed in detail in this study. Names of viral families and genera were indicated with number of species into brackets. For the species selected in the dataset, virus name, abbreviations and accession numbers were indicated. Viral species used as representative member in this study were in bold.

Our focus on sobelivirads (members of the order *Sobelivirales*) enabled the application of modern methodologies to reconstruct their deep evolutionary histories. Notably, the genus *Sobemovirus* exemplifies a unique case where the impact of time-dependent rate phenomenon (TDRP) on underestimation of divergence times using standard molecular clock methods has been considered using the Prisoner of War model (PoW) [29,30]. The PoW model was developed to account for the time-dependent rate phenomenon in viral evolution [31], whereby substitution rate estimates inferred over short timescales tend to decline when extrapolated over deeper evolutionary timescales due to factors like epistasis, nucleotide biases, and substitution saturation. Under this framework, observed genetic divergence is interpreted as arising from a spectrum of effective rate classes : rapidly evolving sites dominate short-term divergence, whereas over longer timescales these sites become increasingly saturated or purged by selection, and more slowly evolving sites contribute proportionally more to the observed divergence. PoW model therefore provides a distance-to-time transformation that maps genetic distances at internal nodes of a tree to calendar time without relying on fixed-node calibration information. In the case of sobelivirads, this framework is especially relevant because standard time-invariant molecular-clock extrapolations using short-term rates underestimate the very ancient origin of sobemoviruses (over four million years) [30,32].

The PoW model application needs three key prerequisites: the detection of a temporal signal in time-stamped sequences, identification of non-recombinant genomic regions, and a robust phylogenetic signal in the data. Here, to overcome the last and more difficult issue at the high taxonomic rank of the order *Sobelivirales*, we leveraged both the structural protein conservation and the advancements in deep-learning algorithms. Indeed, the development of structural phylogenetics, recently accelerated with the efficiency of Alphafold (AF) predictions [33], have demonstrated that 3D data can be used to build more accurate sequence alignments [34] and to reconstruct relationships even between highly divergent proteins in the ‘twilight zone’ of sequence similarity (for review [35]). This approach has been increasingly applied to large collections of viral metagenomic data in order to infer more precise phylogenetic relationships [17,36–38]. By combining it with the discriminant taxonomical criteria for sobelivirads, i.e. polyprotein arrangement, we were able to discard proteins suspected of resulting from evolutionary convergence, to find new candidate barnaviruses from the NCBI Virus database and to select strong and accurate outgroups to reconstruct the sobelivirad phylogeny.

In this paper, we hypothesized that structural modeling would reveal conserved motifs undetectable by sequence alignment alone, and that divergence dates would support ancient horizontal transfers between plant and fungal viruses. We followed an integrative and multi-step approach. Once we had assessed the reliability of the 3D model of RYMV RdRp, we extended our analysis at an intrageneric level revealing the structural conservation among 26 sobemoviral polymerases, including in the divergent sequence domains of the index and thumb. Then, the 3D models of 19 sobelivirad RdRps were generated to produce a structure-guided sequence alignement and to identify the structural features of each viral genera. The RdRp features combined with the characteristic polyprotein arrangement of sobelivirads allowed the identification of new candidate barnaviruses and outgroups. These sequences contributed to generate a new structure-guided alignement and a robust phylogenetic topology. Hence, divergence dates between the two main Sobarna- and Polena-clades and between sobemoviruses infecting plants and barnaviruses infecting fungi were estimated. Finally, molecular signatures of sobelivirads and of each clades were identified refining the definitions of the sobelivirad RdRp conserved motifs. Potential applications of our results including taxonomical revisions, metagenomic sequence assignments, diagnostic tool development and risks of cross-kingdom host jumps were further discussed.

## Results

### Structural features of RYMV RdRp among the picornavirus supergroup

The diversity and evolution of rice yellow mottle virus (*Sobemovirus RYMV*) have been extensively studied [39,40], the estimation of its short-term substitution rate was the starting point of deep evolutionary analyses of sobemoviruses [30,32]. So, we firstly modeled the RYMV RdRp (Fig 1A) using Alphafold v2.2 algorithm [33], which has been shown to predict protein structures with near-experimental accuracy. Predicted 3D model showed local distance difference test (plDDT) scores ranging from 92 to 98, indicating highly reliable structural model (maximum score = 100) (Fig 1A).

**Figure 1:**
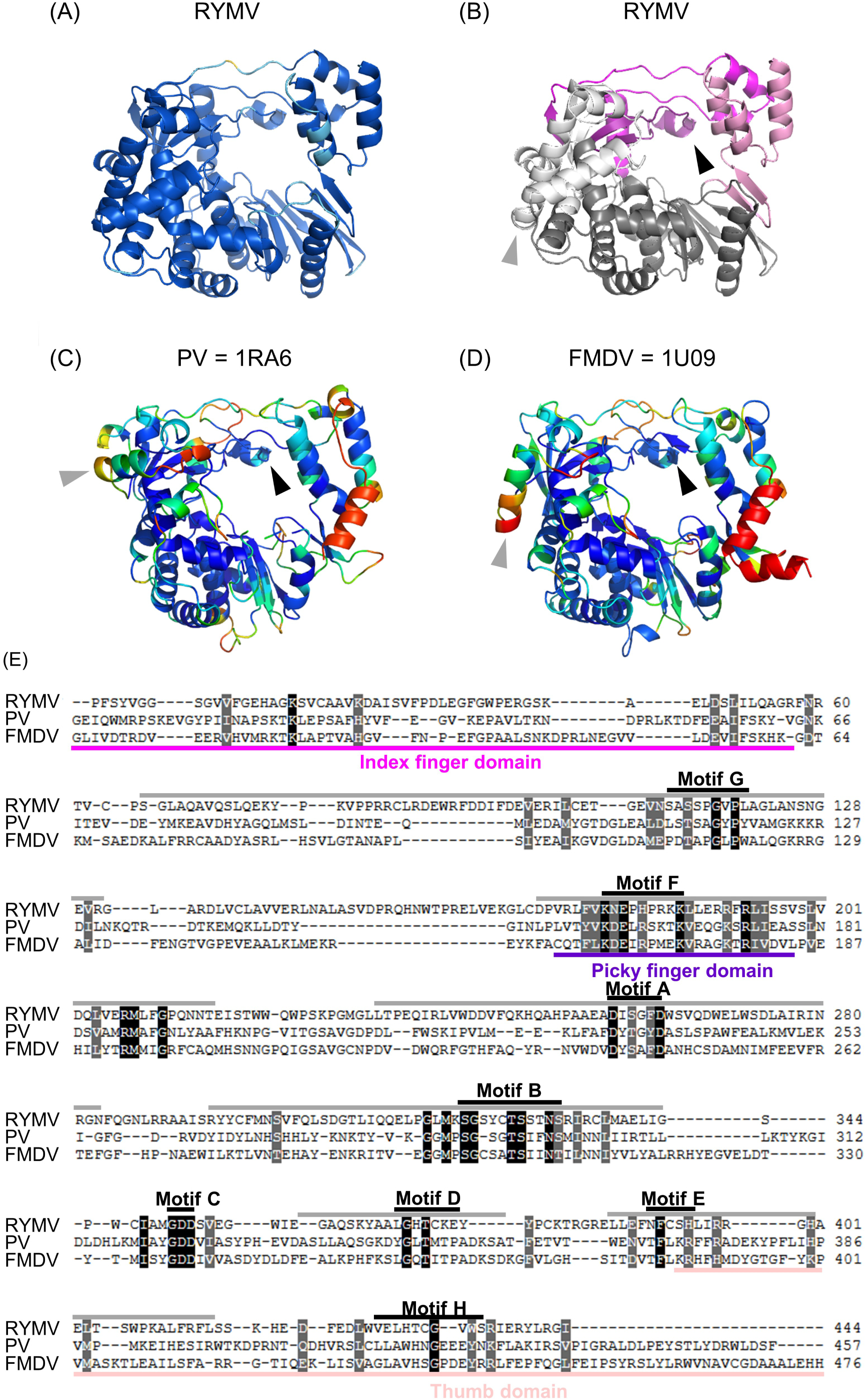
(A) RdRp 3D model of RYMV generated using Alphafold 2.2 colored by per-residue model confidence score (local distance difference test, plDDT). The regions modeled with high confidence (plDDT>90) are shown dark blue. The regions colored light blue (plDDT>70) were modeled with confidence. (B) RYMV RdRp domains. The palm and thumb domains are colored on RYMV model, grey and pink. The index, picky and other fingers are colored magenta, deep purple and white, respectively. (C & D) RdRp structures of picornaviruses (PDB 1RA6, Poliovirus and PDB 1U09, Foot-and-mouth disease virus) are compared to RYMV model and colored based on root mean square deviations (RMSD) to RYMV RdRp model. The regions with high local structural similarity (RMSD<1Å) are shown dark blue. The regions colored light blue (1<RMSD<2) and green (2<RMSD<3) were similar whereas colored other colors, similarity was low. The picornavirus feature, which was the index finger interacted with the top of the thumb trapping the ring finger beneath it, was conserved (black arrow) whereas the α-helix of picornaviruses located at the carboxy-terminal extremity (thumb domain) was lacking in RYMV predicted structure (shown red). The orientation of the pinky finger in the RYMV RdRp model was closer to FMDV rather than PV RdRp structure (grey arrows). Dynamic view and alignment available at https://pat.cbs.cnrs.fr/sobemo/rmsd.html (E) Structure-guided alignment of RdRp sequences of RYMV and the two picornaviruses PV and FMDV using mTM-align. Strictly conserved and similar residues are highlighted (black and grey, respectively), RdRp conserved motifs and homomorphs are outlined (black and grey, respectively). The thumb, index, picky finger domains are outlined pink, magenta and blue violet, respectively.

Up to now, the closest available experimental structures of RdRp belong to poliovirus (PV) and foot-and-mouth disease virus (FMDV) [41], two well-studied species of the order *Picornavirales* in the phylum *Pisuviricota* infecting humans or animals, respectively. The 3D model of RYMV RdRp was aligned with them using mTM-align algorithm [42], a tool that performs global superposition of protein backbones based on TM-scores and RMSD values. The characteristic fingers-palm-thumb organization was observed in all three structures (Fig 1B). Although amino acid conservation between RYMV and PV/FMDV sequences was low (<10% identity) except in the RdRp motifs, the nature and 3D orientation of most secondary structural elements (α-helices and β-sheets) were preserved across all three structures (Fig 1C & 1D). The central part of the protein, which forms the conserved palm domain, showed high local structural similarity, with root mean square deviations (RMSD) below 1 angstrom (Å), validating the approach. One of the major conserved feature of picornavirales RdRps was observed, *i.e.* the index finger at the amino-terminal region is position to interact with the top of the thumb, trapping the ring finger beneath it (Fig 1B). Additionally, the orientation of the so-called pinky finger helix, which has been previously described to distinguish two picornavirad groups including FMDV or PV [41], would bring the predicted structure of RYMV closer to that of FMDV (Fig 1B-D).

The structure-guided alignment generated from the RYMV RdRp model facilitated precise amino acid sequence comparison, in particular for the variable N-terminal domain and the poorly conserved motif H located at the C-terminal extremity (Fig 1E). Globally, only 27 out of 560 sites (4.8%) accross the sequence alignment were strictly identical in the three RdRps. Less than 50% of these conserved residues were located in the ABC motifs, ∼80% in all the 7 RdRp well-known motifs and ∼95% in the entire homomorphs, showing the added value of considering the structural domains rather than the sequence motifs only. However, as expected, the phylogenetic distance between picornavirads and sobelivirads resulted in significant structural differences. In the RdRp core domain, there was a larger number of β-strand elements which are longer in RYMV compared to the two picornaviruses. In contrast, the thumb domain, formed by α-helices at the carboxy-terminal part of RdRps, was smaller in the predicted RYMV structure. Comparison with experimental structures, rare and phylogenetically distant, was informative but insufficient; it must be complemented by modeling of more closely related proteins.

### Structural conservation and phylogeny of sobemoviral RdRps

The RdRp 3D models of the 26 ICTV-approved sobemoviral species (S2 Table) were generated using the same method to assess the structural conservation at the intrageneric scale. They all showed high reliability, with plDDT scores comprised between 90 and 98 (S3 Fig). Three dimensional superpositions of sobemovirus RdRps revealed a high degree of structural similarity in the alpha-carbon traces, with RMSD to RYMV below 1Å (Fig 2A). Only the length of two external loops varied from two to six residues depending the viral species. These loops located at the thumb joint and at the back of the fingers, are named here as « orange and dark green loops », respectively, based on the protein rainbow color code (from N-terminus dark blue to C-terminus red) (Fig 2A).

**Figure 2:**
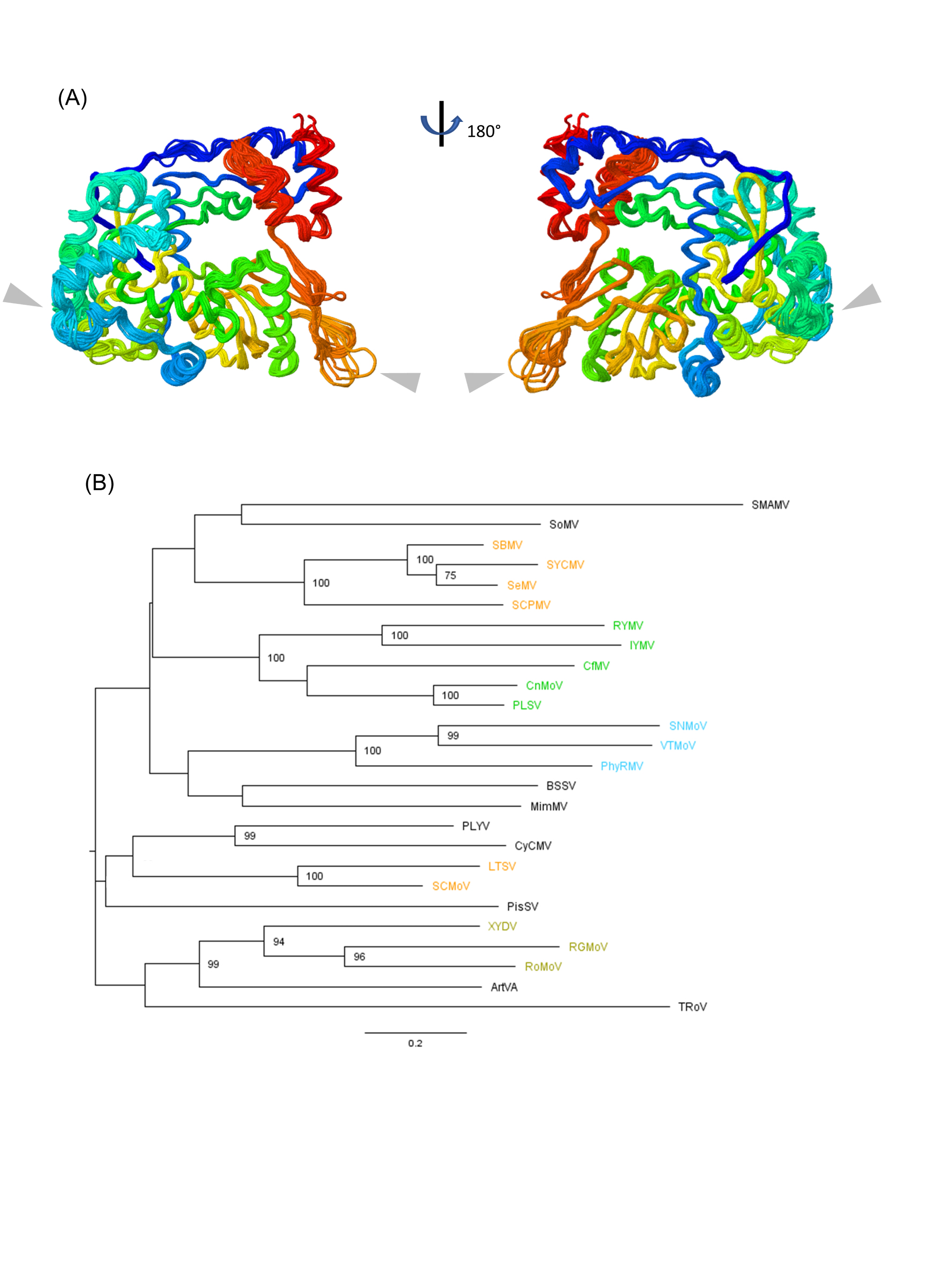
(A) Superimposition of sobemoviral RdRp 3D models (26 species) colored using standard rainbow code (from N-terminus, dark blue, to C-terminus, red). The models showed high reliability (plDDT scores >90) and high local structural similarity (RMSD to RYMV <1Å). Major variable segments at the intra-genera scale are pointed out (grey arrows). Dynamic view and alignment available at https://pat.cbs.cnrs.fr/sobemo/sobemo_v7/sobemo B) Sobemoviral maximum-likelihood phylogenetic tree obtained using the structure-guided mTM-alignment and the GTR+G+I substitution model in SeaView v5.05. Viral species are colored when they belong to lineages with common plant host families as reported in Ghafari et al 2023 (orange, *Fabaceae*, green, *Poaceae,* blue, *Solanaceae* and pale green, *Poales*). Bootstrap supports based on 100 replicates were indicated if up or egal to 75%.

The structure-based alignment of the 26 sobemovirus RdRps computed by mTM-align has a mean length of 446 residues across 472 sites (S4 Fig). With a mean pairwise sequence identity values among sobemovirus RdRps at 54.3% ± 3.2%, the resulting amino acid conservation in the whole mTM-alignment was relatively low, with 120 out of 446 residues being strictly conserved (27% of the mean length). The strictly conserved residues were located in RdRp motifs (31%) or more extended in homomorphs (69%) but 21% belonged to the RdRp extremities upstream and dowstream of the motifs G and D, respectively. Noteworthy, the 75 C-terminal residues including motifs E and H have not been considered in previous sobemoviral alignments (Ghafari et al., 2024). Including this C-terminal part increased alignment length by 18%, and revealed 9 new conserved residues (7.5%).

The RdRp structural alignment was converted to a multiple sequence alignment of nucleic acids by replacing each amino acid by its corresponding codon from their respective viral genomes. The structure-guided alignment was used to reconstruct the maximum-likelihood phylogenetic tree of sobemoviruses using SeaView v5.05 (Fig 2B). Phylogenetic relationships between the viral species confirmed the previously reported lineages and their clustering by plant family hosts [30]. RYMV belong to one of the two lineages infecting monocots host plants of the *Poaceae* family. The other sobemovirus species were distributed in one and two lineages infecting plants of the *Solanaceae* and *Fabaceae* families, respectively, or to different and unique plant families. Hence, the structure-guided alignment of all sobemovirus RdRps recapitulates known phylogenetic relationships, confirming that our approach preserves evolutionary signals even at fine taxonomic scales.

### Structural features and structure-guided alignment of sobelivirad RdRps

The order *Sobelivirales* comprised 122 viral species unevenly distributed into seven genera and three families (Table 1). The genera *Barnavirus, Dinornavirus, Husbsclerovirus* and *Polemovirus* contain only one member recognized by ICTV whereas poleroviruses, sobemoviruses and enamoviruses contain 77, 26 and 15 species, respectively. A subset of 20 RdRps including maximum six taxons in each genera, including sobemoviruses, were selected (Table 1). Given the amino acid sequence similarity between the RdRps of the unique polemovirus and the selected poleroviruses (% ID = 64% ± 0.9%), they were grouped together as « poler/moviruses » for the remainder of the analyses. Indeed, *Polemovirus PNLV* originated from a recombination event between a polerovirus as major parent and a sobemovirus on its CP gene [43].

The RdRp 3D models were generated in a view to guide the sequence alignement at the intergeneric scale. The RdRp models showed plDDT scores ranging mostly from 90 to 98, indicating highly reliable structural models as for the sobemoviral ones (Fig 3A-D and S5 Fig). However, the RdRp 3D model of the Heterocapsa circularisquama RNA virus 01 (HcRNAV), the unique dinornavirus, showed low-confidence plDDT scores (< 60) for 10% of its residues. As a result, dinornaviruses were excluded from dowstream analyses to avoid bias in structural alignments. As expected based on the amino acid sequence conservation (% ID >50%) at the intrageneric scale, comparison of RdRp 3D models confirmed a high structure conservation at this scale (RMSD <1 Å) (Table 2). Sequence and structure conservations of sobemoviral RdRps was significatively lower from those of poler/moviruses.

**Figure 3:**
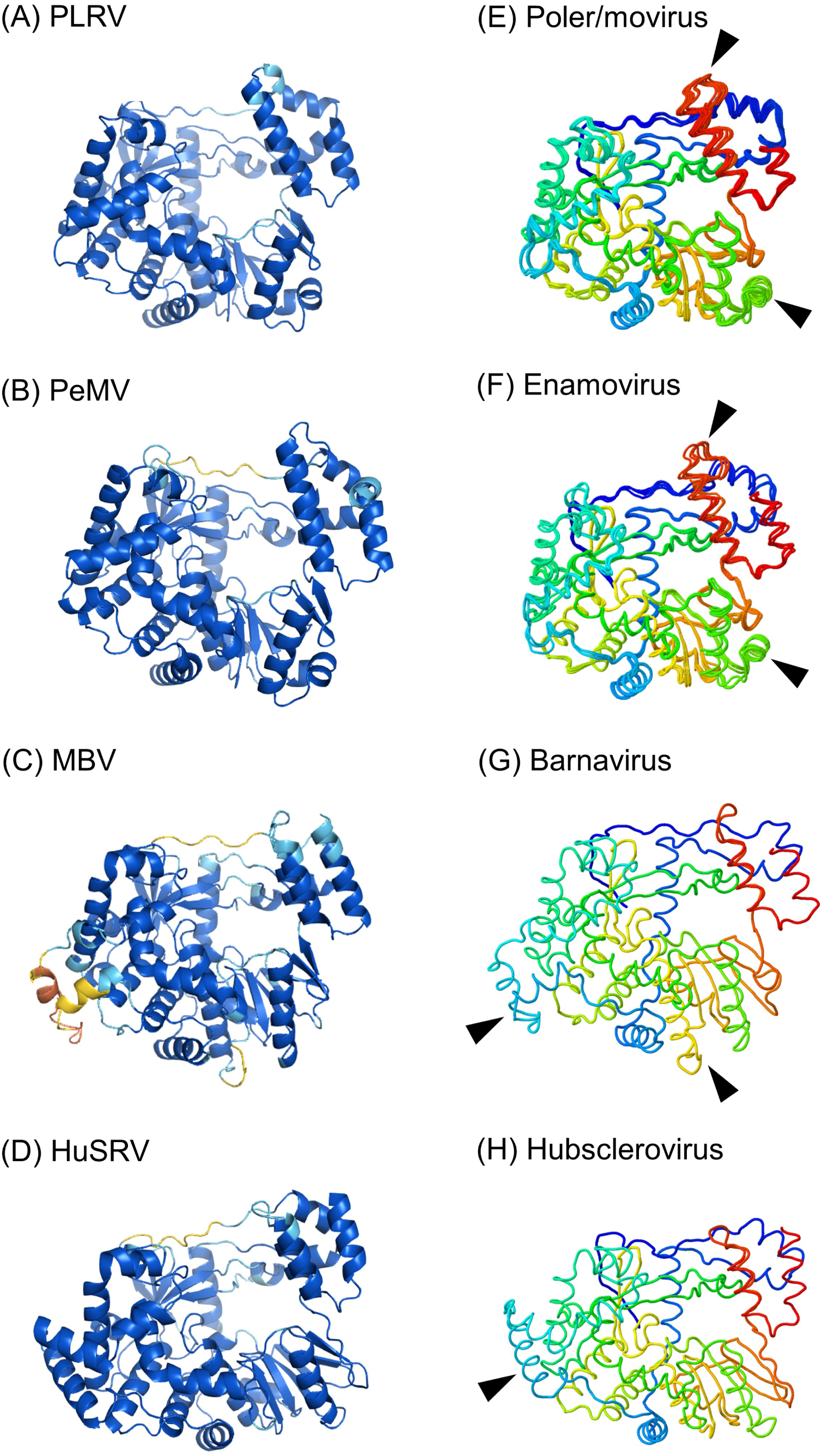
(A-D) RdRp 3D models of genus type members colored by per-residue model confidence score (local distance difference test, plDDT). The regions modeled with high confidence (plDDT>90) are shown dark blue. The regions colored light blue (plDDT>70were modeled with confidence whereas yellow (plDDT>50) and orange (plDDT<50), confidence was low to very low. PLRV (Polerovirus), PeMV (Enamovirus), MBV (Barnavirus), HuSRV (Hubsclerovirus). (E-H) Superimposition of RdRp 3D models by genera colored using standard rainbow code (from N-terminus, dark blue, to C-terminus, red). Poler/movirus (5 species), Enamovirus (4 species), Barnavirus (1 species), Hubsclerovirus (1 species). Structural features at the intergeneric level are pointed out (black arrows see Fig 4). Dynamic view and alignment available at https://pat.cbs.cnrs.fr/sobemo/sobemo_v7/sobeli

**Table 2:** Structure and sequence conservations of sobelivirad RdRps selected in this study.

| Viral genera | Sample number | Structural deviation | Sequence conservation |
| --- | --- | --- | --- |
|  |  | (mean RMSD, Å)* | (mean pairwise % ID)* |
| Sobemovirus | 6 | 0.72 ± 0.12 (a) | 54.3 ± 3.2 (a) |
| Poler/movirus | 7 | 0.56 ± 0.16 (b) | 66.3 ± 4.4 (b) |
| Enamovirus | 4 | 0.62 ± 0.23 (a, b) | 62.8 ± 15.8 (b) |
| Barnavirus | 1 | - | - |
| Hubsclerovirus | 1 | - | - |

\* Letters into brackets (a, b) indicate significative differences between groups (Tukey test, $p < 0.05$ ).
ANOVA : $F(\text{Structure}) = 18.19$ , $p < 0.001$ ; $F(\text{Sequence}) = 10.67$ , $p < 0.001$ .

The 3D superimposition of the 19 RdRp models obtained after structural alignment confirmed the global strutural conservation of sobelivirad RdRps (1.9 Å < RMSD < 2.5 Å) (Fig 4A). While the RdRp core was highly conserved as expected, few additional structural features were observed in some external loops depending on the viral taxons. The « orange loop », found to be highly reliable (plDDT> 80-90) but variable in length among the sobemoviral RdRps, was identified as characteristic of this genera (Fig 2A). By contrast the « dark green loop » was rather stable in most of the sobelivirad RdRps analyzed here, except in those from certain sobemoviruses, and therefore not considered in the dowstream analyses. In the polem/rovirus and enamovirus RdRps, an extra « green helix » was identified at the thumb junction in addition to a longer « red loop » located at the thumb tip. These two specific secondary elements showed a conserved length and high plDDT scores (Fig 3A & 3E). Similarly, high scores were observed for a « light blue helix » located at the edge of the palm in the hubsclerovirus RdRp (Fig 3D & 3H). Interestingly, this extended domain was also observed in the barnavirus RdRp but rather as an unfolded loop (Fig 3C & 3G). However, this « light blue loop » and a « yellow helix » found in the barnavirus RdRp showed low plDDT values (<70) indicating that their 3D conformation remained uncertain. At this step, we cannot exclude that these three structural features were restricted to the unique RdRp sequences of hubsclerovirus and barnavirus species available.

**Figure 4:**
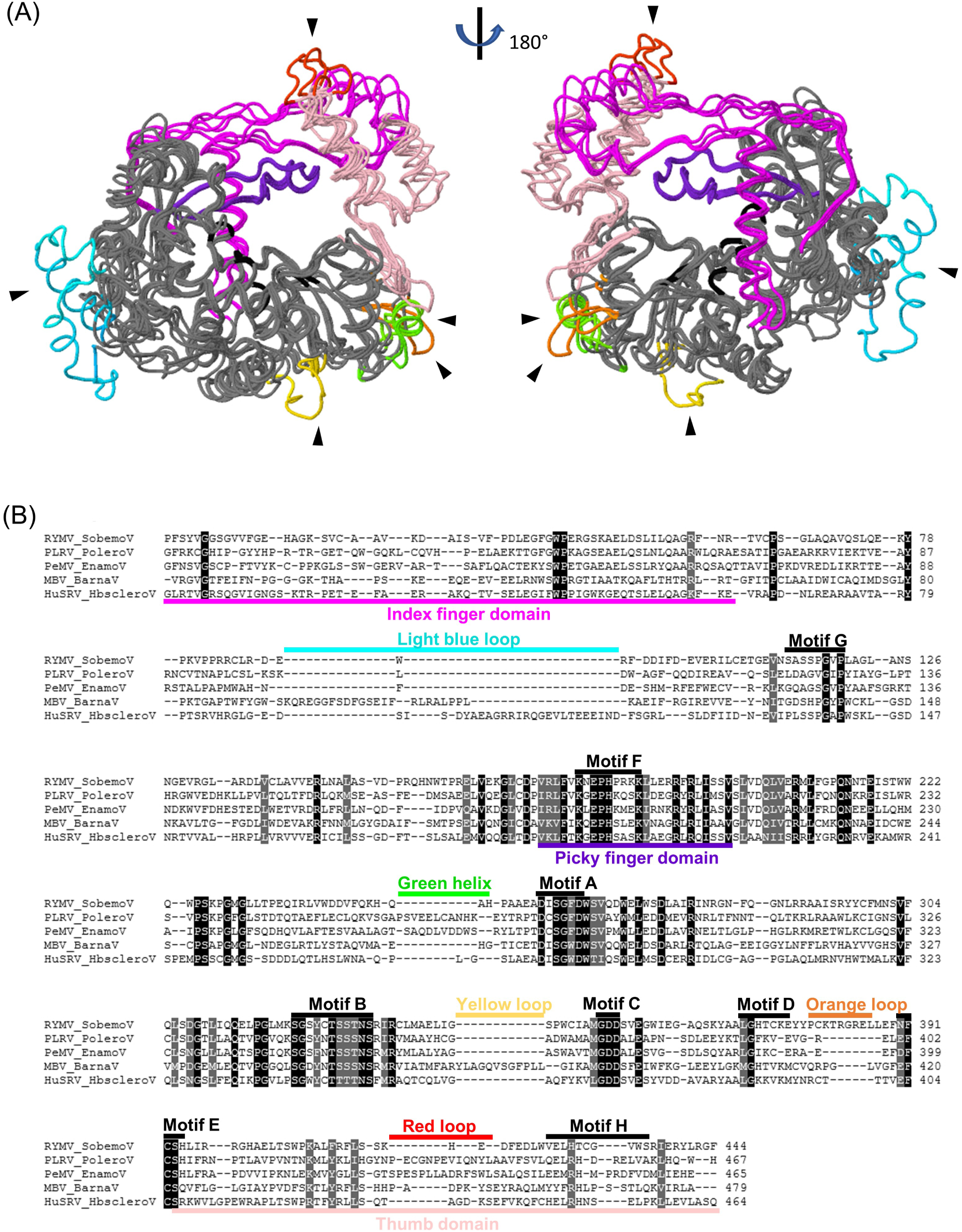
(A) Inter-genera superimposition of RdRp 3D models. For clarity reason, only one model per genera was represented (sobemovirus RYMV, poler/movirus PLRV, enamovirus PeMV, barnavirus MBV, hubsclerovirus HuSRV). The thumb, index, picky finger domains were colored pink, magenta and blue violet, respectively. The conserved motifs A-B-C localized at the center of the channel were colored black. Structural features of the different genera were colored using the rainbow code (as reported in Fig 2 and 3) and pointed out with black arrows. Dynamic view and alignment available at https://pat.cbs.cnrs.fr/sobemo/sobemo_v7/sobeli/ (B) Structure-guided alignment of the 5 sobelivirad RdRp sequences using mTM-align. Strictly conserved and similar residues are highlighted (black and grey, respectively), RdRp conserved motifs are outlined in black. The thumb, index, picky finger domains are outlined pink, magenta and blue violet, respectively. Color code of the structural features of each genera was the same as above. Alignment available at https://pat.cbs.cnrs.fr/sobemo/sobemo_v7/ali19.html

The 3D modeling facilitated sequence alignment and optimization of gap positions and lengths. The structure-guided alignment of the 19 sobelivirad RdRps has a mean length of 456 residues across 581 sites (Fig 4B & S6A-B Fig). Amino acid conservation was notably low, with less than 12% of residues being strictly conserved. As expected, the solvent accessibility scores and predicted secondary structure elements were well conserved along the linear sequences in sobelivirad RdRps. Due to their low sequence conservation, the RdRp extremities are generally not considered in the sequence alignment. However, they play a crucial structural role in the protein structure of picorna-like RdRps (thumb/index finger interaction). The structure-guided approach has enabled us to explicitely include these RdRp extremities in the sequence alignment, for the first time, at this intergeneric taxonomic scale (Fig 4A & 4B).

Intermediate to large insertions and deletions (indels) to account for additional secondary structure elements were heterogeneously distributed along the linear alignment of the RdRps (Fig 4B and S6A-B Fig). They correspond to the secondary structure elements already identified through the global superimposition of the 3D models and could be considered as candidate taxonomical features due to their high intrageneric conservation. For instance, insertions at positions 334-347 and 538-552 were characteristic to the « green helix » and « red loop » in both poler/mo- and enamoviruses and insertions at positions 490-503 to the « orange loop » in sobemoviruses. To a lesser extend due to unique species in the genera and to lower plDDT scores, insertions at positions 120-163 and 443-455 might be characteristic to the « light blue helix or loop » found in the barnavirus or husclerovirus and to the « yellow loop » in the barnavirus. Notably, all these features were located outside the ABC motifs, mostly at the protein extremities. Based on highly reliable structural models, our RdRp sequence alignment contributed to define accurately indel lengths and locations all along the protein sequence and to identify candidate structural features of each sobelivirad genera.

### Identification of new barnaviruses and outgroup RdRp sequences

To balance the underrepresented genera in our RdRps dataset and to confirm the sobelivirad structural features, RdRp sequences of new candidate sobelivirad species were searched in the NCBI Virus database. Ten sequences tagged as husbscleroviruses and polemoviruses were not further considered because they belong to the known HuSRV and PnLV species. By contrast, 69 sequences tagged as classified or unclassified barnavirids were extracted from the database. To avoid false positives due to structural evolutive convergence, we firstly filtered the candidate sequences based on the specific arrangement of domains in the polyprotein (protease / VPg / polymerase) — considered as a major taxonomical criteria of sobelivirads. Thirthy one sequences were identified based on the conserved triad H-D-Gx(S/C)G of the protease and the ABC motif of the polymerase (S7 Table). To avoid redondancy, only one MBV isolate (accession number NC001633) out of three was selected. Based on the high conservation of indels at the intrageneric scale observed in this study, we discarded all the RdRp sequences containing long insertions between the 8 RdRp conserved motifs. Under this stringent filter, five sequences were identified and considered as barnavirus tentative species (Table 3). Three of them, namely CoBV, RsBV and TuBV, were originated from phytoparasitic fungi [44–46], and two, referred here as tBV1 and tBV2, from metagenomic samples [47].

**Table 3:** Virus name, accession numbers and references of the five candidate barnaviruses and the three outgroup viruses used for the phylogenetic analysis of sobelivirads.

| Viral species | Abbr. | NCBI access nb | Uniprot accession nb | Reference |
| --- | --- | --- | --- | --- |
| Colobanthus quitensis associated barnavirus 1 | CoBV | MG686618 | A0A2P1JMX2 | (Nibert et al 2018) |
| Rhizoctonia solani barnavirus 1 | RsBV | KP900904 | A0A0M4KGX8 | (Marzano et al 2016) |
| Tulasnella barnavirus 1 | TuBV | MN738552 | QNN89176 | (Sutella et al 2020) |
| Barnaviridae sp. | tBV1 | MZ218174 | UHS71527 | (Chen et al 2022) |
| Barnaviridae sp. | tBV2 | MZ218185 | UHS71731 | (Chen et al 2022) |
| Soybean thrips sobemo-like virus 5 | SoTV5 | MW023867 | QPZ88400 | [48] |
| Scaphoideus titanus sobemo-like virus 1 | ScTV1 | MN982389 | QIJ56913 | [49] |
| Hypera postica associated sobemovirus 1 | HyPV1 | MW676135 | QUS52861 | [50] |

To reconstruct the evolutionary history of sobelivirad taxons by phylogenetic analyses, a careful outgroup selection is needed. As candidate outgroups must share a common ancestor with sobelivirads, they should also share a similar genomic organisation. Polyprotein sequences tagged as unclassified sobeli-, solemo- and sobemo-like were extracted from the NCBI virus database. From 204 initial sequences, 55 sequences were identified based on the conserved protease and RdRp motifs (S8 Table). Noteworthy, sobemoviruses recently approved by ICTV, such as XYDV and PLSV (accession numbers ON828429 and ON137710), still appeared as unclassified in the NCBI Virus database. After removal of sequences with long insertions and based on basal phylogenetic position, Soybean thrips sobemo-like virus 5 (SoTV5, MW023867), Scaphoideus titanus sobemo-like virus 1 (ScTV1, MN982389), and Hypera postica associated sobemovirus 1 (HyPV1, MW676135) were selected as outgroups for the following analyses (Table 3). They all originated from phytophageous insects [48–50].

The 3D models of the 5 candidate barnaviral and the 3 outgroup RdRps were generated using Alphafold (S9 Fig). They showed plDDT scores ranging mostly from 90 to 98, indicating highly reliable structural models. The low plDDT score of the « light blue loop » of barnavirus RdRp was observed only for MBV but not for the new barnaviruses. The candidate structural features of barnaviruses, the « light blue and yellow loops », showed length variability similarly to the « orange loop » of sobemoviruses. In comparaison to the new barnaviruses, the three outgroup models showed more structural conservation among them. However, the 3D superimposition of the total dataset revealed several specific features in external loops of the 3 outgroups (S10 Fig). Both a longer and a shorter loops were identified in their « dark green » region.

### Sobelivirad phylogeny and divergence date estimation

The structure-guided alignment of the 27 RdRps including candidate barnaviruses and outgroups was generated with mTM-align (Fig 5A & 5B). Following the same method than for the sobemovirus phylogeny, the structural alignment was converted to a multiple sequence alignment of nucleic acids by replacing each amino acid by its corresponding codon, and, then, it was used to construct the maximum-likelihood phylogenetic tree of *Sobelivirales* with SeaView (Fig 6A). The three supplemental sequences were confirmed as outgroups and formed a well-supported cluster. They contributed to reveal two main clades with high bootstrap supports. On the one hand, phylogenetic relationships between poler/moviruses and enamoviruses were confirmed, this clade was referred to as Polena-clade. On the other hand, plant sobemoviruses and fungal barna-/hubsclerovirus clustered together, establishing the monophyly of this group for the first time, referred to as Sobarna-clade. The structure-guided alignment of our relevant dataset resulted in the first accurate phylogeny of sobelivirads — a starting point to propose a taxonomical revision of sobelivirad families.

**Figure 5:**
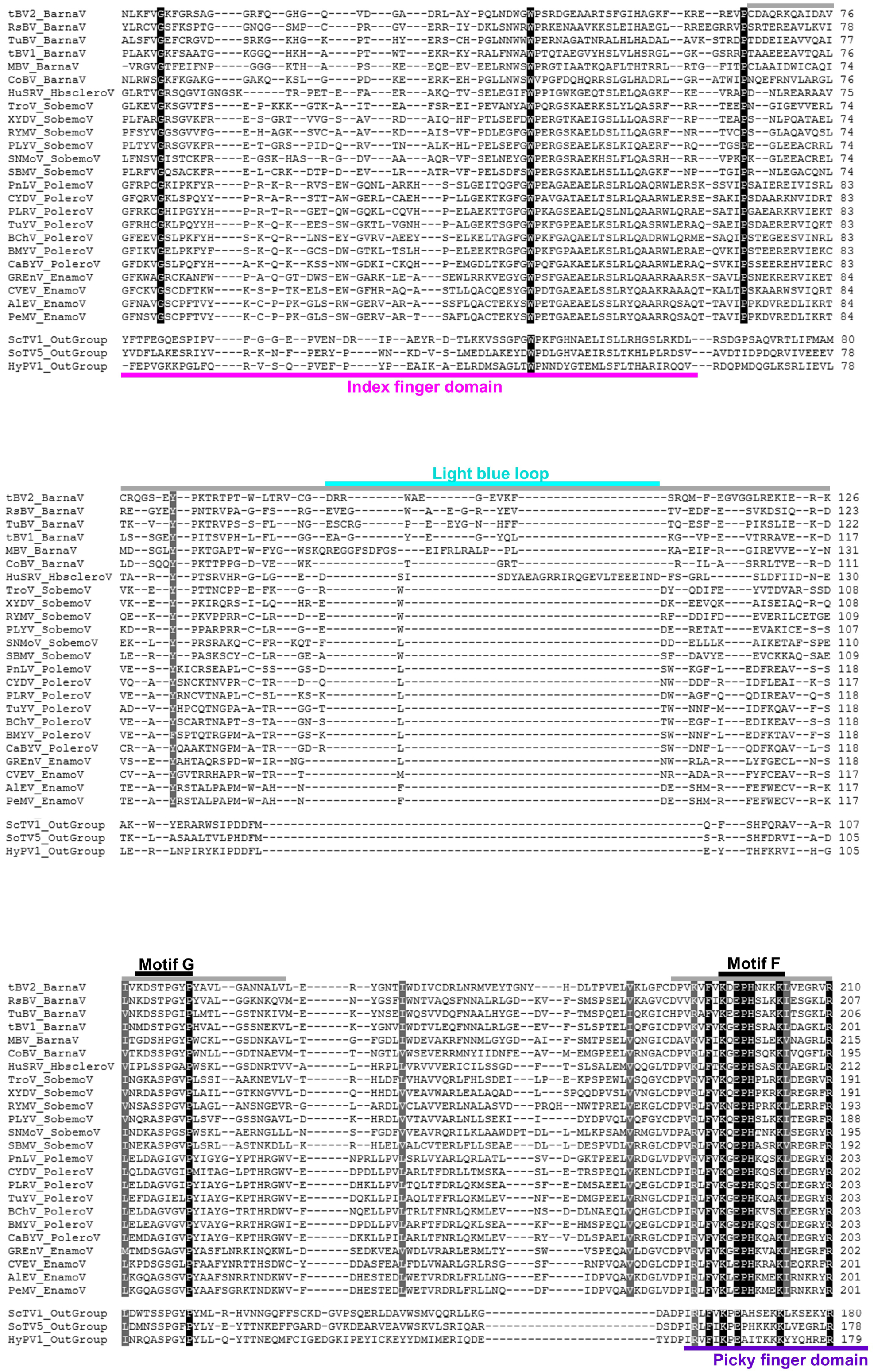
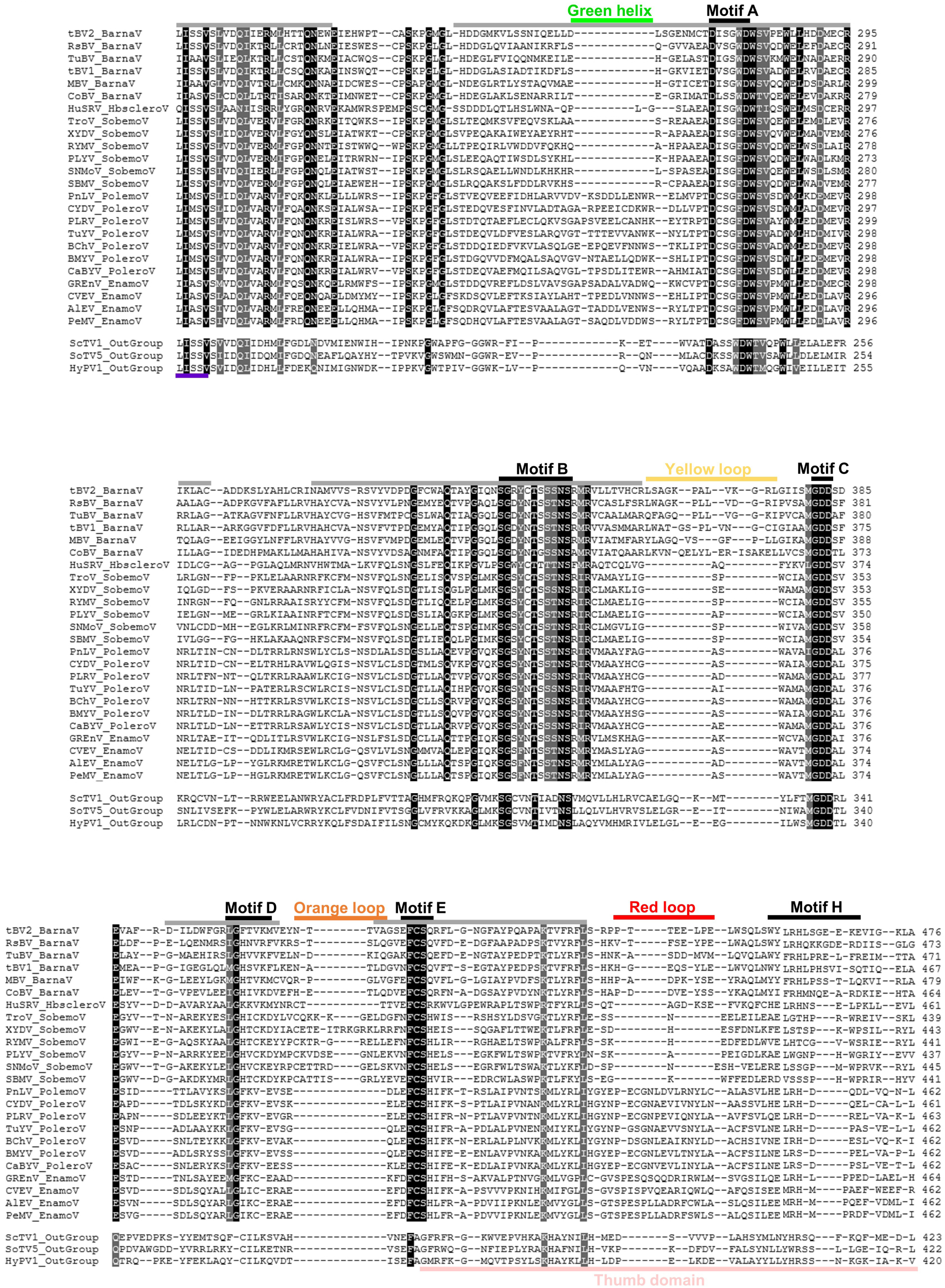
(A-B) Structure-guided alignment of the 27 sobelivirad RdRp sequences using mTm-align. Strictly conserved and similar residues are highlighted (black and grey, respectively), RdRp conserved motifs are outlined in black. The thumb, index, picky finger domains are outlined pink, magenta and blue violet, respectively. Color code of the structural features of each genera was the same as above. Dynamic view and alignment available at https://pat.cbs.cnrs.fr/sobemo/sobemo_v7/sobeli

**Figure 6:**
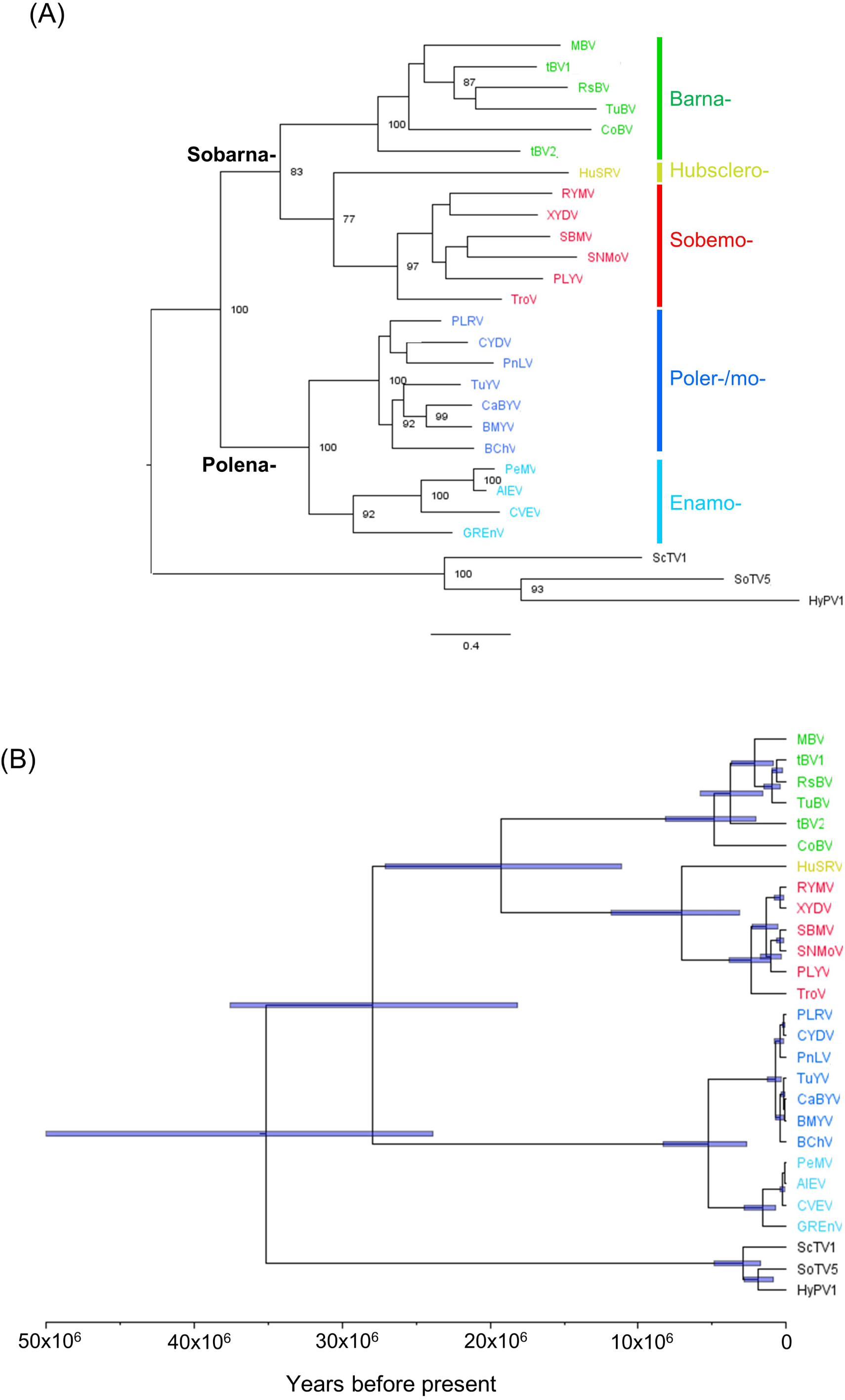
(A) Sobelivirad maximum-likelihood phylogenetic tree obtained using the structure-guided mTm-alignment and the GTR+G+I substitution model in SeaView v5.05. Viral species are colored according to the five genera and the outgroups. Two main clades, namely Sobarna-vs. Polena-clades, were revealed clustering sobemo- and barnaviruses vs. poler/mo- and enamoviruses, respectively. Bootstrap supports based on 100 replicates were indicated if up or egal to 75%. (B) PoW-transformed time tree of the sobelivirads. The horizontal blue bars show 95% highest posterior density of the node age.

To assess the robustness of the inferred topology to the alignment strategy, we also examined an alternative structure-informed alignment based on the 3Di structural alphabet [51,52]. Apart from the position of HuSRV, which appeared distantly related to the sobemoviruses rather than the barnaviruses and the absence of posterior support values, this analysis recovered a broadly similar tree topology (S11 Fig). We therefore retained the mTM-align/codon-based tree for the main analyses, as it enabled downstream nucleotide phylogenetic inference and PoW-based divergence dating. According to this analysis, the most recent common ancestor (TMRCA) of sobemoviruses dates back to 2.2 million years before present (BP) (95% highest posterior density (HPD): 1.0-3.8 million years BP) (Fig 6B), aligning closely with a recent estimate [30]. The divergence time of the Polena-clade was estimated to 5.1 million years BP (95% HPD: 2.6–8.3). The findings here provides the first estimate of the TMRCA for fungal (barna)viruses, which dates back to 4.6 million years BP (95% HPD: 2.0-8.1), and for the Sobarna-clade to 18.7 million years BP (95% HPD: 11.1– 27.1). Furthermore, this analysis offers the first estimation of the TMRCA of the *Sobelivirales* order, reaching back to 27.0 (95% HPD: 18.1-37.5) million years BP, which is nearly ten times older than the origins of sobemoviruses.

### Integrating structural and phylogenetic analyses of sobelivirad RdRps

In this study, 44 sobelivirad polymerases (including 26 sobemoviral RdRps) were modeled and compared. In total, only 43 residues were strictly conserved at the intergeneric scale (less than 9% of the mean sequence length) (Fig 4 & 5). Despite significant divergence in their protein sequences, the RdRp core within the central tunnel of sobelivirads were found to be highly conserved in structure across all genera in agreement with strong functional restraints. A fourth of the conserved residues are located within the ABC motifs, half in the well-characterized RdRp motifs and >80% in homomorphs. Excluding the residues shared with the picornavirus PV and FMDV, 29 out of these 43 conserved amino acids could represent interesting sobelivirad markers for taxonomical assignement, in particular nine of them located in RdRp motifs including 382-SG-383, 288-PH-289, 545-FCS-547 in the motifs A, F and E, respectively (Fig 7).

**Figure 7:**
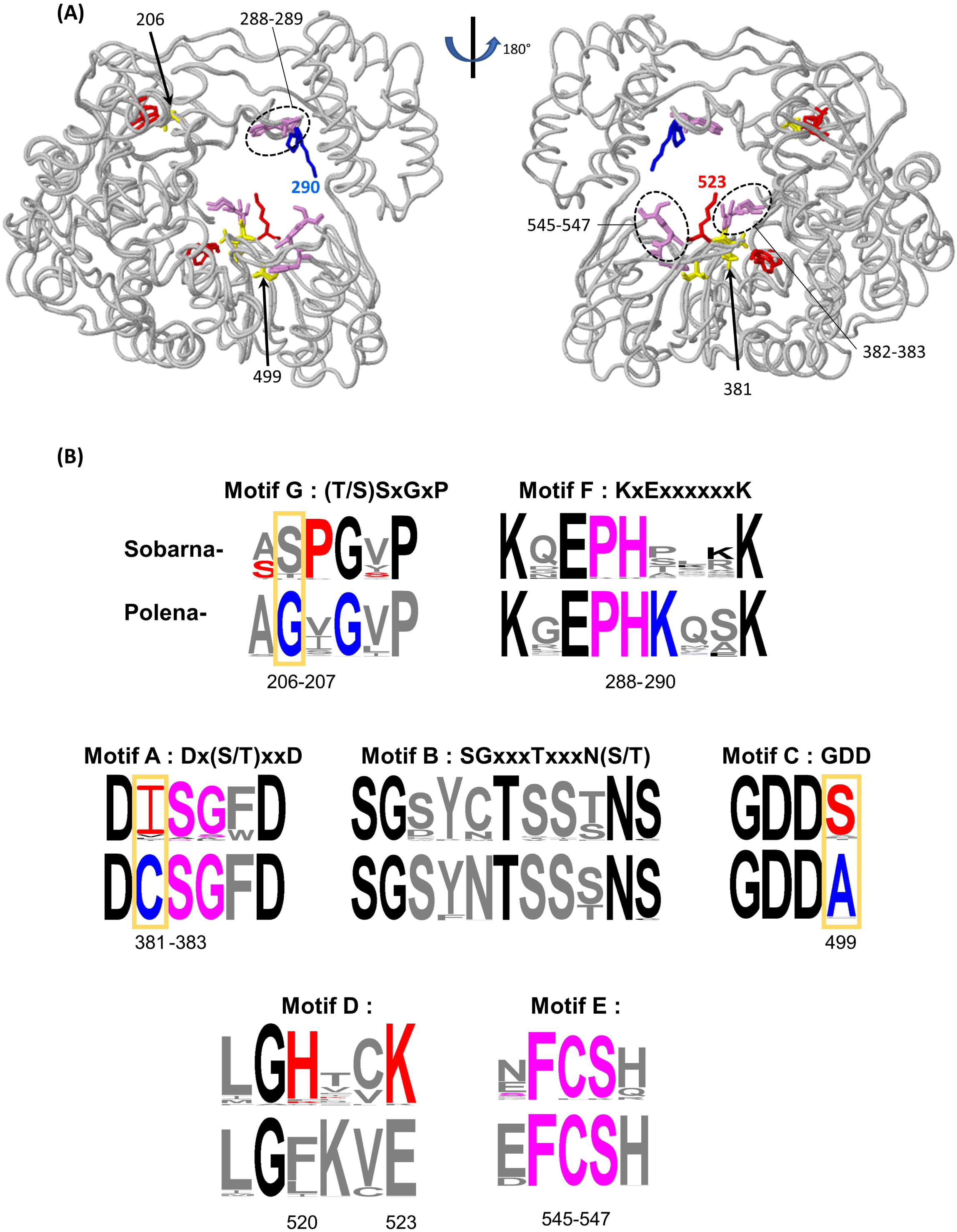
(A) Sobelivirad-specific and clade-specific residues of RdRp motifs located on 3D models. Remarkable residues were figured as sticks on Cα trace of one representative species (sobarna, RYMV and polena-, AlEV, respectively). Sobelivirad-specific, Sobarna- and polena-specific residues were colored magenta, red & blue, respectively, except clade-specific polymorphic positions colored yellow (arrows). Remarkable residues at positions 263, 489 or in interaction 365-486 were indicated by arrows or circled in dotted line (see text). (B) Optimized motifs of sobelivirad RdRps. (a) For each conserved motif, RdRp conserved residues were colored (black), additional sobelivirad-specific residues (magenta), sobarna-specific residues (red) and polena-specific residues (blue). Common clade-specific positions were highlighted (yellow open boxes). Each sequence logo consists of stacks of symbols, one stack for each position in the sequence. The overall height of the stack indicates the sequence conservation at that position, while the height of symbols within the stack indicates the relative frequency of each amino at that position. Position numbering referred to the alignment with 19 sequences.

In order to identify markers of the two major clades identified with the phylogenetic analyses of sobelivirads, the amino acid sequence conservation inside the RdRp core was compared between the Polena- versus Sobarna-clades. All ICTV-recognized poleroviruses and enamoviruses were included in the RdRp sequence dataset in addition to the known sobemoviruses and the candidate barnaviruses (S12 Table). This dataset of 127 sequences allowed the identification of 10 clade-specific residues, *i.e.* amino acids conserved in one clade and distinct in the other clade (Fig 7). Six of them were localized at the same positions 206, 381, and 499 in motifs G, A and C, respectively (referring to the alignment numbering in Fig 5A & 5B). Indeed, the RdRps of the sobarna-clade harbour one isoleucine and two serines at these positions whereas a cysteine, an alanine and a glycine were observed in the polena-clade. Three other residues were specific of the sobarna-clade : 207P in motif G, 520H and 523K, the last two occupying notable 3D positions in motif D. Indeed, The lysine at position 523 pointed into the entry of the central tunnel, while the histidine at position 520 participated to the stabilization of the region with its interaction with the residue 292E (Fig 7). Notably, the sobemoviral « orange loop » between Motifs D and E was also validated in the large dataset. In the Polena-clade, a strictly conserved lysine at position 290 within Motif F on the picky finger domain also pointed into the central tunnel, alongside the conserved sobelivirad lysine at position 293 (Fig 7). In addition, the polena- « green helix » upstream Motif A and the polena- « red loop » dowstream Motif H provided robust indel signatures. Integrating structural and phylogenetic analyses of sobelivirad RdRps revealed molecular signatures of these deeply divergent clades, refining the definitions of the conserved motifs (Fig 7).

## Discussion

Our study aligns with recent efforts to leverage structural phylogenomics for resolving deep evolutionary relationships in highly divergent proteins [35]. We combined AlphaFold, mTM-align and PoW dating to produce a robust and extensive sequence alignement of RdRps and to investigate the evolutionary history of sobelivirads. By contrast to the alignments based on structural alphabets such as FoldSeek 3Di [51], our methodology based on global backbone superpositions offers the advantages to capture extended motifs and to be less sensitive to local inaccuracies in flexible regions. Protein modeling successfully provided high-confidence structural elements to complement anchor points based on the classical functional motifs. Most of these structural features are located outside the ABC motif, often in the RdRp sequence extremities generally not considered for the alignment despite their functional role. For instance, in the NLM’s conserved domain database (CDD) [53], the barnavirid RdRp domain (cd23184) was restricted to the 259 amino acids in the palm domain (corresponding to aa 170-429 in Fig 5A & 5B). However, we showed that the conserved motifs G and H in the index finger and thumb domains were very well predicted by Alphafold (plDDT > 90), and showing high structural similarity (RMSD < 2Å). Hence, our structure-guided approach doubled the alignment length of the sobelivirad RdRp and increased its quality considering now the extremities of RdRp and its intra-molecular interactions. Such an extension can result in a more precise phylogenetic analysis.

As for other mycovirus taxons, barnavirid diversity remains largely unknown. Up to now, the family *Barnaviridae* contains only one viral species recognized by ICTV (*Mushroom bacilliform virus*, MBV). However, the NCBI Virus database contains 69 sequences tagged as « Barnaviridae » and the NCBI Taxonomy browser inventories 13 « unclassified Barnavirus » and 4 « unclassified Barnaviridae ». Moreover, the number of metagenomic publications describing partial sequences related to barnavirids or referred to as « barna-like » viruses is increasing [44–47,54–57]. In the CDD database, besides rhizoctonia solani barnavirus 1 (RsBV) and tulasnella barnavirus 1 (TuBV), apple barna-like 1 (ApBV) is also considered and included in the short protein alignment of the RdRp palm domain. In addition to another independent taxonomical criteria (the polyprotein domain arrangment), our study provides stringent structural criteria of RdRps to assign sequences to the order *Sobelivirales*. We considered that, except the five sequences used in our dataset (RsBV, TuBV, CoBV, tBV1, tBV2), all the other metagenomic sequences cited above should not be considered as barnavirids in absence of other taxonomical evidences. In the near future, new metagenomic datasets targeting fungi could increase the number of barnavirid sequences.

The same approach combining polyprotein domain arrangement and RdRp structural features allowed the identification of three outgroups from metagenomic data based on their genetic distance from the sobelivirad dataset. These outgroups enhanced the quality of the phylogenetic tree and contributed to reveal two main clades with high bootstrap supports. This phylogenic analysis led to a new taxonomic proposal : the creation of two new families Polenaviridae (clustering the genera *Polerovirus*, *Enamovirus* and *Polemovirus*) and Sobarnaviridae (clustering genera *Sobemovirus*, *Barnavirus* and *Hubsclerovirus*), replacing the *Solemoviridae* and *Barnaviridae*. By assessing the conservation of the structure-guided alignment of the 127 RdRp sobelivirid sequences, we enriched the common conserved motifs, identified molecular signatures of the two new proposed families, and revealed structural features that can be used as marker of viral genera. Interestingly, newly identified putative sobemo-like and sobeli-like viruses often represent a significant portion of newly discovered plant and fungus taxa within environnemental viromes [17,47,58–60]. Extending this approach in the near future will accelerate their taxonomic classification, a priority for virus research [61]. Additionally, these RdRp features could be useful for the development of molecular tools for diagnostic and detection such as PCR primers.

At a higher taxonomical rank, in the class *Pisoniviricetes,* the well-known picornaviruses show both structural and functional similarities with sobelivirids, such as the use of a Viral Protein genome-linked (VPg) to prime RNA replication by the RdRp. However, picornaviral VPg are very small (∼20 residues) and are cleaved from the RdRp by a viral protease located downtream of the VPg domain. In contrast, the sobarnavirids and polenavirids share a conserved genomic organization (protease/VPg/polymerase) and have large VPgs (80-150 residues), which likely result in or induce specific molecular mechanisms. While interactions between picornaviral VPg and RdRps and their structure-function relationships have been extensively studied in human and animal viruses, very little data exist for plant viruses. Our approach could help to fill this knowledge gap, or at least provide a better understanding of plant RdRp features through *in silico* comparative analyses.

For example, the comparative analysis of Motif E, unique to RdRps compared to other polymerases, offers a good starting point for better understanding of some of the plant RdRp features. The amino acid composition of this motif is directly linked to the viral replication strategy, whether it is based on protein priming (VPg) or not. Described as a « primer grip », this motif has been shown to help to correctly position the 3’ OH end of the primer strand [9,18]. In picornaviral RdRps, the conserved phenylalanine in Motif E, which faces Motif C, is followed by two residues with relatively long side chains (leucine-lysine, LK), while in flaviviral RdRps, the central channel is narrower, and these residues have rather short side chains (cysteine-serine, CS). Interestingly, plant and fungus sobelivirad RdRps share the same conserved Motif E as flavivirids. Additionaly, the homomorph that contains Motif E forms part of the NTP entry tunnel and can vary in length depending on the viral species. Sobemoviruses exhibit a longer « orange loop » at the thumb joint compared to other members of *Sobelivirales*, another unexpected similarity with flaviviral hepatitis virus C. However, at this taxonomic rank, evolutive convergence should not be discarded.

At the sobelivirad scale, taxonomical factors such as genomic arrangement limit the convergence risk. The common evolutionary history of plant and fungus sobelivirad RdRps could be illustrated by Motif D. This motif influences the rate of nucleotide addition to the nascent RNA strand. Characterized by a glycine hinge and a positively charged lysine, both located in similar positions across available RdRp crystal structures, the distance inside the linear sequence between these two conserved residues varies across different viral families [9]. In the Sobarnaviridae clade, lysine (or its equivalent arginine) is conserved at position N+4, i.e. position 523 (referring to Fig 7). In the Polenaviridae clade, our initial analysis was too stringent to detect the conserved lysine. However, a lysine was found at position N+2 – i.e. position 521, in a large part of the sequences, with its orientation differing from the Sobernaviridae lysine in the 3D structural model. Motif D has been shown to influence not only incorporation speed but also fidelity, both of which are closely linked to viral pathogenesis and virulence. For example, increased fidelity in a poliovirus mutant has been associated with viral attenuation [62]. The putative role of lysine 523 could be tested via site-directed mutagenesis in model sobemoviruses like RYMV.

The structural differences and evolutionary distances between RdRps of sobemoviruses and poleroviruses (two phytovirus genera in the order *Sobelivirales*) were found to be greater than those between sobemoviruses and barnaviruses (a mycovirus genus in the family *Barnaviridae*) as well as hubscleroviruses (a mycovirus genus in the family *Solemoviridae*). Given the role of RdRp as a major marker of viral evolutionary history, our results confirmed that horizontal viral transfert(s) (HVT) between plant/fungal viruses occured in the Order *Sobelivirales*. The direction of the HVT(s) cannot yet be determined without further data on the hosts of metagenomic sequences we used as outgroups.

Using an interdisciplinary approach, we provided the first divergence time estimation between these plant and fungal viruses, dating back to ∼27 million years BP. The TDRP-corrected evolutionary history of *Sobelivirales* was extended five thousand times, far beyond previous estimates [32]. However, this timeframe should be interpreted cautiously. Our current application of the PoW model is based on a short-term substitution rate estimated for the RYMV polymerase gene and assumes that this rate provides a reasonable proxy across the broader sobelivirad phylogeny. If short-term rates vary substantially among plant- and fungal-infecting clades, some node ages may be shifted to older or younger than their true values. More generally, while PoW accounts for time-dependent rate decay across timescales, it does not fully resolve lineage-specific branch-rate heterogeneity. These estimates should therefore be viewed as a first unified temporal framework for sobelivirad evolution rather than as precise point estimates for all clades. Nevertheless, even with our current conservative estimates, the estimate time favour a model in which the present-day distribution of sobelivirads across plants and fungi reflects past host shifts across kingdoms rather than uninterrupted codivergence with their current hosts over hundreds of millions of years [63]. Other lineages of mycoviruses either with single- or double-stranded RNA genomes also showed close phylogenetic relationships with plant viruses at different taxonomical ranks [20,24,64], suggesting that taxonomy of these viruses should not be considered independently but rather as a continuum. An interdisciplinary approach at the interface of biophysics, bioinformatics, and virology should be encouraged to better understand evolution, interactions and biological mechanisms of myco- and phytoviruses.

## Material and methods

### Sequence datasets

The sobemoviral dataset used in this study contained all the RdRp sequences from RYMV to the other 25 ICTV-approved sobemoviral species (S2 Table). The sobelivirad dataset contained a subset of representative RdRp sequences from the viral order (Table 1). Six poleroviruses, six sobemoviruses and four enamoviruses were selected to maximize the RdRp sequence diversity for each genera based on their phylogeny. For each of the genera *Barnavirus, Husbsclerovirus* and *Polemovirus,* only one ICTV-approved member was available. The unique member of *Dinornavirus* (HcRNAV) was excluded from the dataset due to low-confidence plDDT scores (see above).

Five putative barnaviral sequences were added to the 19 sobelivirad datased based on a three-step methodology. Firstly, polyprotein sequences tagged as « Barnaviridae » or « unclassified Barnaviridae » were extracted from the NCBI Virus database and aligned with MUSCLE [65] in SeaView v5.05 [66]. The N-terminal extremities of the RdRps often incomplete were manually refine. Secondly, the presence of the conserved triad H-D-Gx(S/C)G of the protease upstream the ABC motif of the polymerase were checked (S7 Table). Finally, sequences with long insertions between the 8 RdRp conserved motifs were discarded (Table 3). In addition, three outgroup sequences tagged as unclassified sobeli-, solemo- and sobemo-like were selected and added to result in the 27 RdRp dataset using the same process (S8 Table, Table 3). A total sobelivirad dataset made of 127 RdRp sequences was also constitued adding all ICTV-recognized poleroviruses and enamoviruses (S12 Table) to the 26 sobemoviral and the 27 sobelivirad datasets described above.

### 3D model prediction, evaluation and structural alignment

The 3D structures of the RYMV, the 25 other sobemoviral and the 27 sobelivirad RdRps were predicted using a locally installed version of Alphafold v2.2 [33]. For each sequence, 5 models were generated using default Alphafold parameters and the highest-ranked model was conserved. The pLDDT reliability scores were available in the B-factor field of each Alphafold model. Models showing 10% of their residues having low-confidence plDDT scores (< 60) were not considered in this study.

The RYMV 3D model was superposed to the closest experimental structures available, PDB 1RA6 and 1U09 of poliovirus (PV) and foot-and-mouth disease virus (FMDV) RdRps, respectively, using mTM-align [42]. Pairwise RMSD scores were calculated using ProFit (Martin A.C.R., http://www.bioinf.org.uk/software/profit/). The aligned sequences and their corresponding RMSD scores in Angstroms (identifier suffixed with d3.RMSD_1ra6A an .d3.RMSD_1u09A) are interfaced with the 3D models using Jsmol applet and script [67] (https://pat.cbs.cnrs.fr/sobemo/rmsd.html). The Cα deviation colors can be projected onto one of the displayed structures by cliking one of these identifiers. The pairwise sequence identity between the proteins was calculated with Ident and Sim software [68]. Descriptive statistics (mean and standard deviation), one-way analysis of variance (ANOVA) and Tukey’s honest significant difference (HSD) post-hoc test were performed to identify specific pairwise differences (p < 0.05). The strictly conserved and the similar residues were visualized onto the sequence alignment using Color Align Conservation [68]. Homomorphs were identifed based on [18].

The 3D models of the 26 sobemoviral and the 27 sobelivirad RdRps were superposed using mTM-align and were interfaced with their sequences using JSmol applet and script (https://pat.cbs.cnrs.fr/sobemo/sobemo_v7/sobemo and https://pat.cbs.cnrs.fr/sobemo/sobemo_v7/sobeli, respectively). By default, the 3D models were colored using standard rainbow code (from N-terminus, dark blue, to C-terminus, red). Selected Alphafold models can be displayed and customized by using the toggle buttons and the popup menus from the right bottom panel and the colors of the amino acids of one aligned sequence from the top panel can be projected on the selected 3D structure(s) by clicking on the corresponding protein identifier in the top left panel. We carefully delineated the polymerase domain shared by the proteins by inspecting both the positional conservation of the aligned sequences, the superposition of their predicted secondary structures and the solvent accessibility. The secondary structure and the solvent accessibility of each sobelivirad model residue was calculated using STRIDE [69].

### Phylogenetic analysis

The mTM-align structural alignments of sobemoviral and sobelivirad RdRps were translated into multiple sequence alignments of nucleic acids by replacing each amino acid by their corresponding codon from the viral genomes using PAL2NAL [70]. JModelTest 2 [71] was used to identify the best nucleotide substitution model for phylogenetic analysis based on the Akaike Information Criterion. The Maximum-Likelihood tree was then constructed with the General Time Reversible Model with an optimised rate variation and class for invariant sites (GTR + G + I) using PhyML implemented in SeaView v 5.05 [66]. The reliability of the phylogenetic tree nodes was estimated via a bootstrap resampling procedure with 100 replicates. The sobemoviral and sobelivirad phylogenetic tree were midpoint rooted and rooted using the outgroup branch, respectively. The mTM-align tree was compared to an alternative one based on the 3Di structural alphabet using Foldtree powered by Foldseek [51,52].

### Time-tree reconstruction using the Prisoner of War model

We used the Prisoner of War (PoW) model of virus evolution to account for the power-law decay in the substitution rate estimates of sobelivirads, as described in Ghafari et al. [30], where we also established the presence of a robust and representative molecular clock signal for the RdRp locus for sobemoviruses. Briefly, the short-term substitution rate for the RYMV polymerase gene, inferred from a standard molecular clock model was used as the baseline rate. RYMV provides the only available sobelivirad lineage for which a robust short-term rate estimate had previously been established. PoW model was then applied to root-to-node genetic distances in an ultrametric distance tree. For each internal node, the genetic node height was transformed into calendar time using the PoW distance-to-time mapping, parameterised by the short-term rate distribution estimated from RYMV and the fastest-evolving rate class. Calendar-time branch durations were then obtained as the differences between transformed parent and child node ages. The maximum clade credibility tree was constructed after 100 iterations using TreeAnnotator v.1.10.4 and visualized through a lineages-through-time plot using the phytools library in R [72].

### Polymerase evolution

The 44 sequences of sobemoviral and sobelivirad RdRps were compared alone and with those of PV and FDMV using Color Align Conservation to identify candidate sobelivirad markers. Then, the total sobelivirad dataset made of 127 RdRp proteins were aligned using MAFFT [73] to identify RdRp core markers of the Polena- versus Sobarna- clades. The three outgroups were not considered in this analysis. At each site of the 8 RdRp conserved motifs, amino acids present in all sequences of one clade (or that were similar based on BLOSUM score ≥ 0) and absent from sequences of the other clade were considered as clade-specific residue. The location of these sobelivirad and clade-specific marker residues were visualized and colored on the 3D models of RYMV and PLRV as sticks on Cα trace using JSmol. Motif visualization was performed with WebLogo v2.8.2 [74]. Each sequence logo consists of stacks of symbols, one stack for each position in the sequence. The overall height of the stack indicates the sequence conservation at that position, while the height of symbols within the stack indicates the relative frequency of each amino at that position.

## Acknowledgements

This work benefited from discussions with Aurore Comte and Nils Poulicard, other members of Virus/Cereal Interactions in Tropical Agroecosystems group and colleagues of Plant Health Institute Montpellier. This work was supported by the French Infrastructure for Integrated Structural Biology (FRISBI) ANR-10-INBS-0005 and a Wellcome Trust Early-Career Award (grant 309205/Z/24/Z) to M.G.

## Data availability

The sequence accession numbers used in this study and their alignments were given in Tables, Figures and Supporting data. The 3D models generated using Alphafold were available online.

## Supporting data

**S1 Table :** Taxonomy and biological characteristics in the phylum Pisuviricota.

**S2 Table :** Dataset of the 26 sobemoviral sequences.

**S3 Figure** : RdRp 3D models of sobemoviruses colored by per-residue model confidence score (local distance difference test, plDDT).

**S4 Figure** : Structure-guided alignment of RdRp sequences of the 26 sobemoviruses using mTm-align.

**S5 Figure** : RdRp 3D model of pole/moviruses, enamoviruses and dinornavirus colored by per-residue model confidence score (local distance difference test, plDDT).

**S6 Figure** : (A-B) Structure-guided alignment of RdRp sequences of the 19 sobelivirads using mTm-align.

**S7 Table** : Dataset of 31 sequences from NCBI Virus database, tagged as Barnaviridae and containing both protease and sobelivirad RdRp conserved motifs.

**S8 Table** : Dataset of 55 sequences from NCBI Virus database, tagged as unclassified sobeli-, solemo- and sobemo-like and containing both protease and RdRp conserved motifs

**S9 Figure** : (A-B) RdRp 3D models of candidate barnaviruses and outgroups colored by per-residue model confidence score (local distance difference test, plDDT). (C-D) Superimposition of RdRp 3D models of barnaviruses and outgroups.

**S10 Figure** : Structure-guided alignment of RdRp sequences of the 6 barnaviruses, 3 outgroups and 4 sobelivirads representative of each other genera.

**S11 Figure :** Phylogenetic tree obtained using Foldtree.

**S12 Table** : Additionnal 99 sequences (genus, name species and accession numbers).

