## Supplementary material for "Integrating structural modeling and divergence dating of RNA-dependent RNA polymerases to resolve the evolutionary history of plant and fungal viruses: from sobemoviruses to sobelivirads": S1 Table

**S1 Table :** Taxonomy and biological characteristics in the phylum *Pisuviricota*

| Class | Order | Suborder | Family | Genome | Virion morphology | Number of particles | Number of segments | Genome size (bp) | Host |
|---|---|---|---|---|---|---|---|---|---|
| *Pisoniviricetes* | *Picornavirales* | | *Picornaviridae* | +ssRNA | icosahedral | 1 | 1 | 6.7-10.1 | vertebrate |
| | | | *Caliciviridae* | +ssRNA | icosahedral | 1 | 1 | 7.4-8.3 | vertebrate |
| | | | *Dicistroviridae* | +ssRNA | icosahedral | 1 | 1 | 8-10 | invertebrate |
| | | | *Iflaviridae* | +ssRNA | icosahedral | 1 | 1 | 9-11 | invertebrate |
| | | | *Marnaviridae* | +ssRNA | icosahedral | 1 | 1 | 8.6-9.6 | protists |
| | | | *Noraviridae* | +ssRNA | icosahedral | 1 | 1 | 12.3 | invertebrate |
| | | | *Polycipiviridae* | +ssRNA | icosahedral | 1 | 1 | 10-12 | invertebrate |
| | | | *Secoviridae* | +ssRNA | icosahedral | 1-2 | 1-2 | 9-13.7 | plants |
| | | | *Solinviviridae* | +ssRNA | icosahedral | 1 | 1 | 10-11 | invertebrate |
| | ***Sobelivirales*** | | ***Solemoviridae*** | +ssRNA | icosahedral | 1 | 1 | 4-6 | **plants** |
| | | | ***Barnaviridae*** | +ssRNA | bacilliform | 1 | 1 | 4 | **fungi** |
| | | | *Alvernaviridae* | +ssRNA | icosahedral | 1 | 1 | 4.4 | protists |
| | *Nidovirales* | *Abnidovirineae* | *Abyssoviridae* | +ssRNA | ? | | 1 | 35.9-36 | invertebrate |
| | | *Arnidovirineae* | *Arteviridae* | +ssRNA | spherical | 1 | 1 | 12.7-15.7 | invertebrate, vertebrate |
| | | | *Cremegaviridae* | +ssRNA | spherical | 1 | 1 | 17 | vertebrate |
| | | | *Gresnaviridae* | +ssRNA | spherical | 1 | 1 | 18 | vertebrate |
| | | | *Olifoviridae* | +ssRNA | spherical | 1 | 1 | 15 | vertebrate |
| | | *Cornidovirineae* | *Coronaviridae* | +ssRNA | spherical | 1 | 1 | 22-36 | vertebrate |
| | | *Mesnidovirineae* | *Medionviridae* | +ssRNA | sherical | 1 | 1 | 20.2-25 | invertebrate |
| | | | *Mesoniviridae* | +ssRNA | spherical | 1 | 1 | 20 | invertebrate |
| | | *Monidovirineae* | *Mononiviridae* | +ssRNA | spherical | 1 | 1 | 41.1 | invertebrate |
| | | *Nanidovirineae* | *Nanghoshaviridae* | +ssRNA | spherical | 1 | 1 | 13 | vertebrate |
| | | | *Nanhypoviridae* | +ssRNA | spherical | 1 | 1 | 18 | vertebrate |
| | | *Ronidovirineae* | *Euroniviridae* | +ssRNA | bacilliform | 1 | 1 | 24.6-29.3 | invertebrate |
| | | | *Roniviridae* | +ssRNA | bacilliform | 1 | 1 | 26-29 | invertebrate |
| | | *Tornidovirineae* | *Tobaniviridae* | +ssRNA | spherical | 1 | 1 | 28 | vertebrate |
| *Duplopiviricetes* | *Durnavirales* | | *Amalgaviridae* | dsRNA | helical | 1 | 1 | 3.5 | plants |
| | | | *Curvulaviridae* | dsRNA | icosahedral | 1 | 2 | 1.7-2.3 | fungi |
| | | | *Fusariviridae* | dsRNA | capsidless | - | 1 | 5.9-10.7 | fungi |
| | | | *Hypoviridae* | +ssRNA | capsidless | - | 1 | 7.3-18.3 | fungi |
| | | | *Partitiviridae* | dsRNA | icosahedral | 2 | 2 | 3-4.8 | fungi, plants, protists |
| | | | *Picobirnaviridae* | dsRNA | icosahedral | 1 | 2 | 1.7-1.9 & 2.4-2.7 | invertebrate, vertebrate |
| | | | *Soropartitiviridae* | dsRNA | ? | 1 | 2 | 1.6-1.7 & 1.6-1.9 | bacteria |
| *Stelpaviricetes* | *Patatavirales* | | *Potyviridae* | +ssRNA | helical | 1-2 | 1-2 | 8.2-11.5 | plants |
| | *Stellavirales* | | *Astroviridae* | +ssRNA | icosahedral | 1 | 1 | 6.8-7 | vertebrate |
| *Unassigned* | *Yadokarivirales* | | *Yadokariviridae* | +ssRNA | trans-encapsided | 1 | 1 | 2.8-6.4 | fungi |
| | *Unassigned* | | *Hadakaviridae* | +ssRNA | capsidless | - | 10-11 | 14.3-15.3 | fungi |
