## Supplementary material for "Integrating structural modeling and divergence dating of RNA-dependent RNA polymerases to resolve the evolutionary history of plant and fungal viruses: from sobemoviruses to sobelivirads": S2 Table

**S2 Table :** Dataset of the 26 sobemoviral sequences

| Virus name | Abbr. | accession number |
|---|---|---|
| Artemisia virus A | ArtVA | NC017914 |
| Blueberry shoestring virus | BSSV | LC081344 |
| Cocksfoot mottle virus | CfMV | DQ680848 |
| Cymbidium chlorotic mosaic virus | CyCMV | LC381945 |
| Cynosorus mottle virus | CnMoV | OM323994 |
| Imperata yellow mottle virus | IYMV | NC011536 |
| Lucerne transient streak virus | LTSV | NC001696 |
| Mimosa mosaic virus | MimMV | OP456085 |
| Papaya lethal yellowing virus | PLYV | NC018449 |
| Physalis rugose mosaic virus | PhyRMV | MK681145 |
| Pistacia sobemovirus | PisSV | MT334602 |
| Poaceae Liege sobemovirus | PLSV | ON137710 |
| Rice yellow mottle virus | RYMV | AJ608207 |
| Rottboellia yellow mottle virus | RoMoV | KC577469 |
| Ryegrass mottle virus | RGMoV | EF091714 |
| Sesbania mosaic virus | SeMV | NC002568 |
| Snake melon asteroid mosaic virus | SMAMV | MT989351 |
| Solanum nodiflorum mottle virus | SNMoV | NC033706 |
| Southern bean mosaic virus | SBMV | AF055887 |
| Southern cowpea mosaic virus | SCPMV | NC001625 |
| Sowbane mosaic virus | SoMV | GQ845002 |
| Soybean yellow common mosaic virus | SYCMV | KX096577 |
| Subterranean clover mottle virus | SCMoV | AY376451 |
| Turnip rosette virus | TRoV | NC004553 |
| Velvet tobacco mottle virus | VTMoV | NC014509 |
| Xufa yellow dwarf virus | XYDV | ON828429 |
