## Supplementary material for "Integrating structural modeling and divergence dating of RNA-dependent RNA polymerases to resolve the evolutionary history of plant and fungal viruses: from sobemoviruses to sobelivirads": S3 Fig

**S3 Figure :** RdRp 3D models of sobemoviruses colored by per-residue model confidence score (local distance difference test, plDDT).

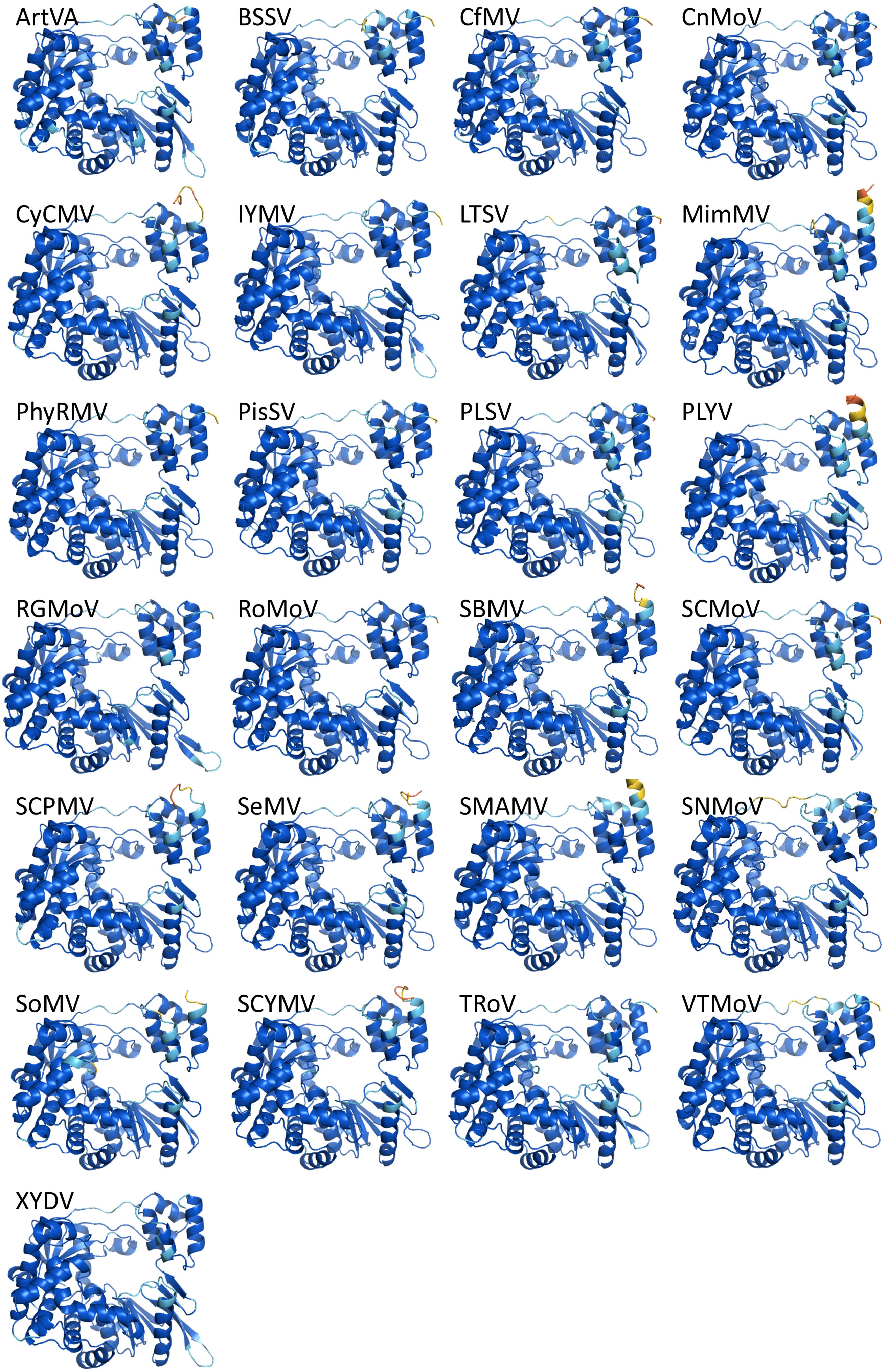
