## Supplementary material for "Integrating structural modeling and divergence dating of RNA-dependent RNA polymerases to resolve the evolutionary history of plant and fungal viruses: from sobemoviruses to sobelivirads": S5 Fig

**S5 Figure :** RdRp 3D model of poleroviruses (BChV, BMYV, CYDV, CaBYV, TuYV), enamoviruses (AEV, CVEV, GEV), polemovirus (PnLV) and dinornavirus (HcRNAV) colored by per-residue model confidence score (local distance difference test, plDDT). The regions modeled with high confidence (plDDT>90) are shown dark blue. The regions colored light blue (plDDT>70) were modeled with confidence whereas yellow (plDDT>50), confidence was low.

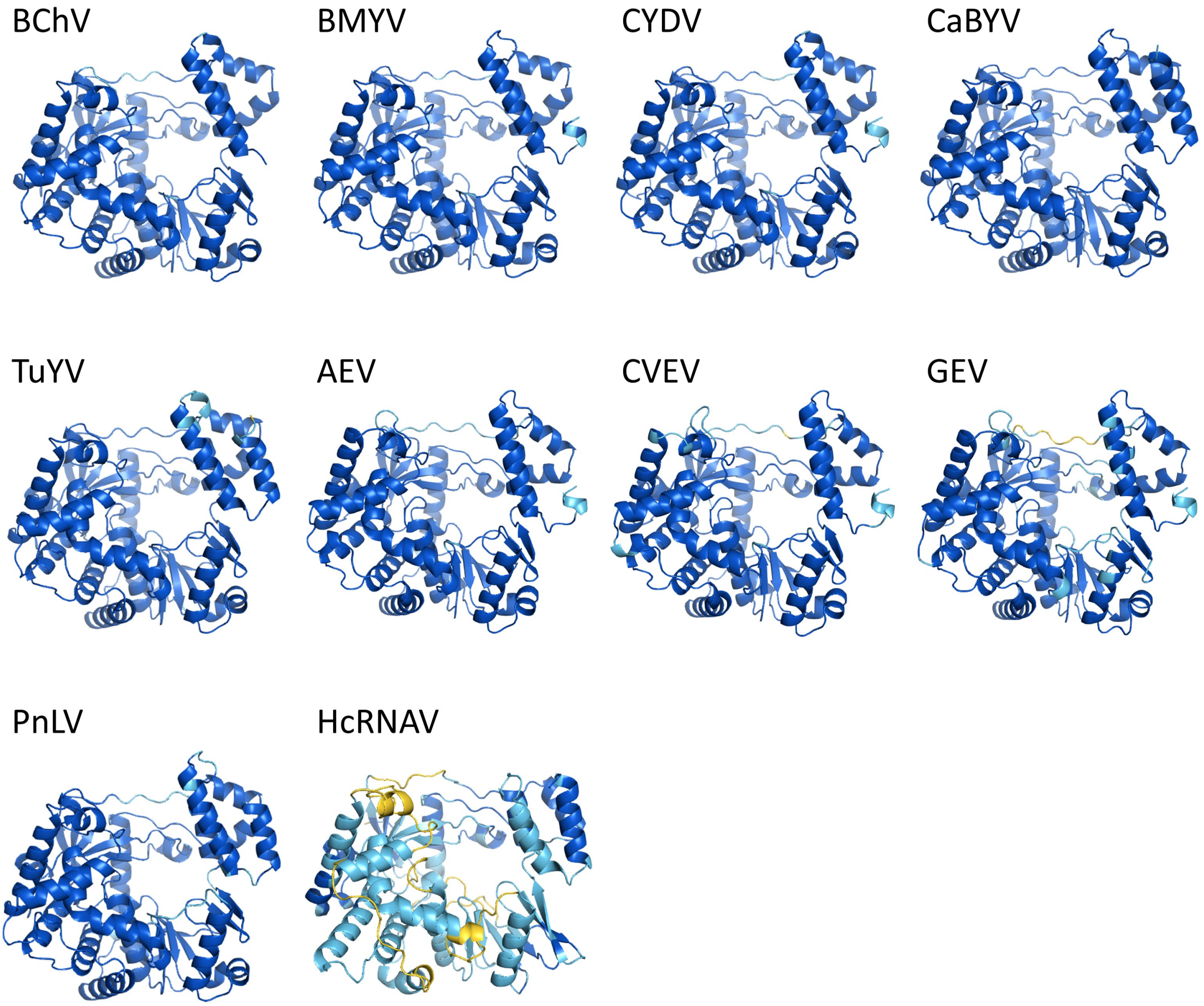
