## Supplementary material for "Integrating structural modeling and divergence dating of RNA-dependent RNA polymerases to resolve the evolutionary history of plant and fungal viruses: from sobemoviruses to sobelivirads": S6A Fig

**S6A Figure :** Structure-guided alignment of RdRp sequences of the 19 sobelivirads using mTm-align. Strictly conserved and similar residues are highlighted (black and grey, respectively), RdRp conserved motifs and homomorphs are outlined (black and grey, respectively). The thumb, index, picky finger domains are outlined pink, magenta and blue violet, respectively.

Color Align Conservation results

```
MBV_BarnaV        -VRGVGTFEIFNP-----GGG--KTH--A--PS--KE--EQE-EV-EELRNWSWPRGTIAATKQAFLTHTRRL--RTG-FITPCLAAIDWICAQIMDSGL 79
HuSRV_HbscleroV   GLRTVGRSQGVIGNGSK------TR-PETE-FA--ER--AKQ-TV-SELEGIFWPPIGWKGEQTSLELQAGKF--KE--VRAPD--NLREARAAVTAR-- 78
TroV_SobemoV      GLKEVGKSGVTFS----E-P-KKK--GTKA-IT--EA--FSR-EI-PEVANYAWPQRGSKAERKSLYLQASRF--RR--TEEPN--GIGEVVERLVKE-- 77
XYDV_SobemoV      PLFARGRSGVKFR----E-S-GRT--VVGS-AV--RD--AIQ-HF-PTLSEFDWPERGTKAEIGSLLLQASRF--RP--TEAPS--NLPQATAELVKE-- 77
RYMV_SobemoV      PFSYVGGSGVVFG----E-H-AGK--SVCA-AV--KD--AIS-VF-PDLEGFGWPERGSKAELDSLILQAGRF--NR--TVCPS--GLAQAVQSLQEK-- 77
PLYV_SobemoV      PLTYVGRSGVKFR----E-T-GRS--PIDQ-RV--TN--ALK-HL-PELSEFGWPERGSEAELKSLKIQAERF--RQ--TGSPE--GLEEACRRLKDR-- 77
SNMoV_SobemoV     LFNSVGISTCKFR----E-GSK-HAS--RG-DV--AA--AQR-VF-SELNEYGWPERGSRAEKHSLFLQASRH--RR--VPKPK--GLEEACRELEKL-- 77
SBMV_SobemoV      PLRFVGQSACKFR----E-L-CRK--DTPD-EV--LR--ATR-VF-PELSDFSWPERGSKAELHSLLLQAGKF--NP--TGIPR--NLEGACQNLLER-- 77
PnLV_PolemoV      GFRPCGKIPKFYR----P-R-K-R--RVSEWGQNLARKH--S-SLGEITQGFGWPEAGAEAELRSLRLQAQRWLERSKSSVIPSAIEREIVISRLVES-- 86
CYDV_PoleroV      GFQRVGKLSPQYY----P-R-A-R--STTAWGERLCAEH--P-LLGEKTKGFGWPAVGATAELTSLRLQAARWLERSESAKIPSDAARKNVIDRTVQA-- 86
PLRV_PoleroV      GFRKCGHIPGYYH----P-R-T-R--GETQWGQKLCQVH--P-ELAEKTTGFGWPKAGSEAELQSLNLQAARWLQRAESATIPGAEARKRVIEKTVEA-- 86
TuYV_PoleroV      GFRHCGKLPQYYH----P-K-Q-K--EESSWGKTLVGNH--P-ALGEKTSGFGWPKFGPEAELKSLRLQASRWLERAQSAEIPSDAERERVIQKTADV-- 86
BChV_PoleroV      GFEEVGSLPKFYH----S-K-Q-R--LNSEYGVRVAEEY--S-ELKELTAGFGWPKFGAQAELTSLRLQADRWLQRMESAQIPSTEGEESVINRLVEA-- 86
BMYV_PoleroV      GFIKVGELPKFYF----S-K-Q-K--GCSDWGTKLTSLH--P-ELEEKTRGFGWPKFGPAAELKSLRLQAARWLERAEQVKIPSTEERERVIEKCVEA-- 86
CaBYV_PoleroV     GFDKVGSLPQFYH----A-K-Q-K--KSSNWGDKICKQH--P-EMGDLTKGFGWPQFGAKAELKSLRLQAARWLERAQSVKIPSSEEREHVIERCCRA-- 86
GREnV_EnamoV      GFKWAGRCKANFW----P-A-Q-GT-DWSEWGARKLE-A--SEWLRRKVEGYGWPSFGAEAELRSLRLQAARRAARSKSAVLPSNEKRERVIKETVSE-- 87
CVEV_EnamoV       GFCKVGSCDFTKW----K-S-P-TK-ELSEWGFHRAQ-A--STLLQACQESYGWPDTGAEAELSSLRYQAAKRAAAQTKALTPSKAARWSVIQRTCVA-- 87
AlEV_EnamoV       GFNAVGSCPFTVY----K-C-P-PK-GLSRWGERVAR-A--SSFLQACTEKYSWPETGAEAELSSLRYQAARRQSAQTTAVIPPKDVREDLIKRTTEA-- 87
PeMV_EnamoV       GFNSVGSCPFTVY----K-C-P-PK-GLSSWGERVAR-T--SAFLQACTEKYSWPETGAEAELSSLRYQAARRQSAQTTAVIPPKDVREDLIKRTTEA-- 87
```

Index finger domain

Light blue loop

Motif G

```
MBV_BarnaV        Y--PKTGAPTWFYG-WSKQREGGFSDFGS--EIFRLRALP-PL---------------------KAEIF-RGIREVVE--Y-NITGDSHPGYPWCKL--G 146
HuSRV_HbscleroV   Y--PTSRVHRGLG--E--D----------SI---------SDYAEAGRRIRQGEVLTEEEIND-FSGRL---SLDFIID-N-EVIPLSSPGAPWSKL--G 145
TroV_SobemoV      Y--PTTNCPPEFK--G--R----------W---------------------------------DY-QDIFE-YVTDVAR-SSDINGKASPGVPLSSI--A 123
XYDV_SobemoV      Y--PKIRQRSILQ--HR-E----------W---------------------------------DK-EEVQK-AISEIAQ-R-QVNRDASPGVPLAIL--G 123
RYMV_SobemoV      Y--PKVPPRRCLR--D--E----------W---------------------------------RF-DDIFD-EVERILCETGEVNSASSPGVPLAGL--A 124
PLYV_SobemoV      Y--PPARPRRCLR--G--E----------W---------------------------------DE-RETAT-EVAKICE-S-SVNQRASPGVPLSVF--G 122
SNMoV_SobemoV     Y--PRSRAKQCFR-KQT-F----------L---------------------------------DD-ELLLK-AIKETAF-SPEINDKASPGSPWSKL--A 125
SBMV_SobemoV      Y--PASKSCYCLR--GE-A----------W---------------------------------SF-DAVYE-EVCKKAQ-SAEINEKASPGVPLSRL--A 124
PnLV_PolemoV      YKICRSEAPLCSS--GS-D----------L---------------------------------SW-KGF-LEDFREAV--S-SLELDAGIGVPYIGYG-Y 134
CYDV_PoleroV      YSNCKTNVPRCTR--D--Q----------L---------------------------------NW-DDF-RIDFLEAI--K-SLQLDAGVGIPMITAG-L 133
PLRV_PoleroV      YRNCVTNAPLCSL--KS-K----------L---------------------------------DW-AGF-QQDIREAV--Q-SLELDAGVGIPYIAYG-L 134
TuYV_PoleroV      YHPCQTNGPAATR--GG-T----------L---------------------------------TW-NNF-MIDFKQAV--F-SLEFDAGIELPYIAYG-K 134
BChV_PoleroV      YSCARTNAPTSTA--GN-S----------L---------------------------------TW-EGF-IEDIKEAV--S-SLELDAGVGVPYIAYG-T 134
BMYV_PoleroV      FSPTQTRGPMATR--GS-K----------L---------------------------------SW-NNF-LEDFKTAV--F-SLELEAGVGVPYVAYG-R 134
CaBYV_PoleroV     YQAAKTNGPMATR--GD-R----------L---------------------------------SW-DNF-LQDFKQAV--L-SLEMDAGIGVPYIAYG-K 134
GREnV_EnamoV      YAHTAQRSPDWIR-NG-------------L---------------------------------NW-RLA-RLYFGECL--N-SMTMDSGAGVPYASFLNR 135
CVEV_EnamoV       YGVTRRHAPRWTR--T-------------M---------------------------------NR-ADA-RFYFCEAV--R-SLKPDSGSGLPFAAFYNR 134
AlEV_EnamoV       YRSTALPAPMWAH--N-------------F---------------------------------DE-SHM-RFEFWECV--R-KLKGQAGSGVPYAAFSNR 134
PeMV_EnamoV       YRSTALPAPMWAH--N-------------F---------------------------------DE-SHM-RFEFWECV--R-KLKGQAGSGVPYAAFSGR 134
```

Motif F

```
MBV_BarnaV        SDNKAVLTG--FGDLIWDEVAKRFNNMLGYGD-AI--F-SMTPSELVQNGICDAVKVFIKQEPHSLEKVNAGRLRIIAAVGLVDQIVTRLLCMKQNNAEI 240
HuSRV_HbscleroV   SDNRTVVAL--HRPLLVRVVVERICILSSGD--F---T-SLSALEMVQQGLTDPVKLFTKGEPHSASKLAEGRLRQISSVSLAANIISRRLYGRQNRVEK 237
TroV_SobemoV      AKNEVLVTR--HLDFLVHAVVQRLFHLSDEI--LP--E-KPSPEWLVSQGYCDPVRVFVKQEPHPLRKLDEGRVRLISSVSLVDQLVERVLFGRQNRKEI 216
XYDV_SobemoV      TKNGVVLDQ--HHDLVVEAVWARLEALAQAD--L---SPQQDPVSLVVNGLCDPVRLFVKQEPHPKRKLREGRFRLISSVSLIDQLVERVLFGYQNSLEI 216
RYMV_SobemoV      NSNGEVRGL--ARDLVCLAVVERLNALASVD--PRQH--NWTPRELVEKGLCDPVRLFVKNEPHPRKKLLERRFRLISSVSLVDQLVERMLFGPQNNTEI 218
PLYV_SobemoV      SSNGAVLDK--HRDLVVTAVVARLNLLSEKI--I-----DYDPVQLVQFGYCDPVRLFVKQEPHSLKKITEGRFRLISSVSLVDQLVERMIFGPQNELEI 213
SNMoV_SobemoV     ERNGLLLNS--FGDFVVEAVRQRILKLAAWDPTDL--L-MLKPSAMVRMGLVDPARVFVKQEPHTNKKLSEGRYRLISSVSIVDQIIERLLFGPQNQLEI 220
SBMV_SobemoV      STNKDLLKR--HLELVALCVTERLFLLSEAE--DL--L-DESPVDLVRRGLCDPVRLFVKQEPHASRKVREGRFRLISSVSLVDQLVERMLFGPQNQLEI 217
PnLV_PolemoV      PTHRGWVENPRLLPVLSRLVYARLQRLATL---SV--D-GKTPEELVRDGLVDPVRVFVKGEPHKQSKLDEGRYRLIMSVSLIDQLVARVLFQKQNKLEL 228
CYDV_PoleroV      PTHRGWVEDPDLLPVLARLTFDRLLTMSKA---SL--E-TRSPEQLVKENLCDPIRLFVKQEPHKQSKLDEGRYRLIMSVSLIDQLVARVLFQAQNKSEI 227
PLRV_PoleroV      PTHRGWVEDHKLLPVLTQLTFDRLQKMSEA---SF--E-DMSAEELVQEGLCDPIRLFVKGEPHKQSKLDEGRYRLIMSVSLVDQLVARVLFQNQNKREI 228
TuYV_PoleroV      PTHRGWVEDQKLLPILAQLTFFRLQKMLEV---NF--E-DMGPEELVRNGLCDPIRLFVKGEPHKQAKLDEGRYRLIMSVSLVDQLVARVLFQNQNKREI 228
BChV_PoleroV      RTHRDWVFNQELLPVLTRLTFNRLQKMLEV---NS--D-DLNAEQLVQHGLCDPIRVFVKGEPHKVSKLEEGRYRLIMSVSLVDQLVARVLFQNQNKREI 228
BMYV_PoleroV      RTHRGWIEDPDLLPVLARFTFDRLQKLSEA---KF--E-HMSPEQLVQEGLCDPIRLFVKGEPHKQSKLDEGRYRLIMSVSLVDQLVARVLFQNQNKREI 228
CaBYV_PoleroV     PTHRGWVEDKKLLPILARLTFNRLQKMLEV---RY--V-DLSPAELVRRGLCDPIRVFVKGEPHKQAKLDEGRYRLIMSVSLVDQLVARVLFQNQNKREI 228
GREnV_EnamoV      KINQKWLDSEDKVEAVWDLVRARLERMLTY---SW-----VSPEQAVLDGVCDPVRVFVKGEPHKVAKLHEGRFRIIASVSMVDQLVARMLFQSQNKQEL 227
CVEV_EnamoV       TTHSDWCYDDASFEALFDLVWARLGRLRSG---SF-----RNPVQAVQDGLCDPIRLFVKLEPHKRAKIEQKRFRLIASVSLADQLVARMLFQEQNQAEL 226
AlEV_EnamoV       RTNDKWVFDHESTEDLWETVRDRLFRLLNG---EF-----IDPVQAVKDGLVDPIRLFVKLEPHKMEKIRNKRYRLIASVSIVDQLVARMLFREQNEEEL 226
PeMV_EnamoV       KTNDKWVFDHESTEDLWETVRDRLFRLLNQ---DF-----IDPVQAVKDGLVDPIRLFVKLEPHKMEKIRNKRYRLIASVSIVDQLVARMLFRDQNEEEL 226
```

Picky finger domain
