## Supplementary material for "Integrating structural modeling and divergence dating of RNA-dependent RNA polymerases to resolve the evolutionary history of plant and fungal viruses: from sobemoviruses to sobelivirads": S7 Table

**S7 Table :** Dataset of 31 sequences from NCBI Virus database, tagged as Barnaviridae and containing both protease and sobelivirad RdRp conserved motifs

| Virus name | Accession number |
| --- | --- |
| Apple barna-like virus 1 | MN386956 |
| Colobanthus quitensis associated barnavirus 1 | MG686618 |
| Mushroom bacilliform virus | NC001633 |
| Mushroom bacilliform virus | U07551 |
| Mushroom bacilliform virus | KY357511 |
| Pleurotus pulmonarius barnavirus 1 | PV579973 |
| Pleurotus sapidus barnavirus 1 | PV579972 |
| Rhizoctonia solani barnavirus 1 | KP900904 |
| Sclerotinia sclerotiorum barnavirus 1 | MT646362 |
| Sonisav virus | PP173898 |
| Tulasnella barnavirus 1 | MN738552 |
| Barnaviridae sp. | MW826417 |
| Barnaviridae sp. | MZ218171 |
| Barnaviridae sp. | MZ218174 |
| Barnaviridae sp. | MZ218178 |
| Barnaviridae sp. | MZ218179 |
| Barnaviridae sp. | MZ218180 |
| Barnaviridae sp. | MZ218182 |
| Barnaviridae sp. | MZ218185 |
| Barnaviridae sp. | MZ218186 |
| Barnaviridae sp. | MZ218190 |
| Barnaviridae sp. | MZ218191 |
| Barnaviridae sp. | MZ218193 |
| Barnaviridae sp. | MZ218199 |
| Barnaviridae sp. | MZ218202 |
| Barnaviridae sp. | MZ218205 |
| Barnaviridae sp. | MZ218206 |
| Barnaviridae sp. | MZ218208 |
| Barnaviridae sp. | MZ218210 |
| Barnaviridae sp. | MZ218213 |
| Barnaviridae sp. | MZ218216 |
