## Supplementary material for "Integrating structural modeling and divergence dating of RNA-dependent RNA polymerases to resolve the evolutionary history of plant and fungal viruses: from sobemoviruses to sobelivirads": S8 Table

**S8 Table** : Dataset of 55 sequences from NCBI Virus database, tagged as unclassified sobeli-, solemo- and sobemo-like and containing both protease and RdRp conserved motifs

| Virus name | Accession number |
|---|---|
| Frankliniella occidentalis associated sobemo-like virus 2 | MN725051 |
| Hubei sclerotinia RNA virus 1 | MT646396 |
| Hubei sclerotinia RNA virus 1 | MK889164 |
| Hypera postica associated sobemovirus 1 | MW676135 |
| Laodelphax striatellus sobeli-like virus 2 | LC851051 |
| Laodelphax striatellus sobeli-like virus 4 | LC851053 |
| Mortimer virus | MN167486 |
| Mute swan feces associated sobemovirus 1 | MW588173 |
| Mute swan feces associated sobemovirus 1 | MW588174 |
| Poaceae Liege sobemovirus | ON137710 |
| Reticulitermes chinensis sobeli-like virus 1 | BK067197 |
| Reticulitermes flavipes sobeli-like virus 2 | BK067200 |
| Reticulitermes speratus sobeli-like virus 1 | BK067201 |
| Ripiglev virus | PP173612 |
| Sea buckthorn enamo-like virus | BK068687 |
| Scaphoideus titanus sobemo-like virus 1 | MN982389 |
| Sichuan mosquito sobemo-like virus | MZ556266 |
| Sopihev virus | PP173919 |
| Soybean thrips sobemo-like virus 5 | MW023867 |
| Stellaria aquatica mottle polerovirus A | OP389993 |
| Xufa yellow dwarf virus | ON828429 |
| Sobemovirus sp | MZ218224 |
| Sobemovirus sp | MZ218227 |
| Sobemovirus sp | MZ218228 |
| Sobemovirus sp | MZ218229 |
| Sobemovirus sp | MZ218230 |
| Sobemovirus sp | MZ218231 |
| Sobemovirus sp | MZ218232 |
| Sobemovirus sp | MZ218234 |
| Sobemovirus sp | MZ218235 |
| Sobemovirus sp | MZ218239 |
| Sobemovirus sp | MZ218241 |
| Sobemovirus sp | MZ218242 |
| Sobemovirus sp | MZ218243 |
| Sobemovirus sp | MZ218244 |
| Sobemovirus sp | MZ218245 |
| Solemoviridae sp | OR871272 |
| Solemoviridae sp | OR871275 |
| Solemoviridae sp | MZ395980 |
| Solemoviridae sp | MW826523 |
| Solemoviridae sp | MW826560 |
| Sobelivirales sp | OR843431 |
| Sobelivirales sp | OR843435 |
| Sobelivirales sp | OR843437 |
| Sobelivirales sp | OR843442 |

| | |
|---|---|
| Sobelivirales sp | OR843444 |
| Sobelivirales sp | OR843456 |
| Sobelivirales sp | OR843456 |
| Sobelivirales sp | OR843461 |
| Sobelivirales sp | OR843463 |
| Sobelivirales sp | OR843476 |
| Sobelivirales sp | OR843477 |
| Sobelivirales sp | OR843478 |
| Sobelivirales sp | OR843479 |
| Sobelivirales sp | OR843480 |
