## Supplementary material for "Integrating structural modeling and divergence dating of RNA-dependent RNA polymerases to resolve the evolutionary history of plant and fungal viruses: from sobemoviruses to sobelivirads": S9 Fig

**S9 Figure :** (A-B) RdRp 3D models of candidate barnaviruses and outgroups colored by per-residue model confidence score (local distance difference test, plDDT). The regions modeled with high confidence (plDDT>90) are shown dark blue. The regions colored light blue (plDDT>70) were modeled with confidence whereas yellow (plDDT>50), confidence was low. (C-D) Superimposition of RdRp 3D models of barnaviruses (one ICTV-approved and the 5 tentative species) and outgroups (3 species) colored using standard rainbow code (from N-terminus, dark blue, to C-terminus, red). Structural features at the intergeneric level are pointed out (black arrows see Fig 4). Dynamic view and alignment available at https://pat.cbs.cnrs.fr/sobemo/sobemo_v7/sobeli

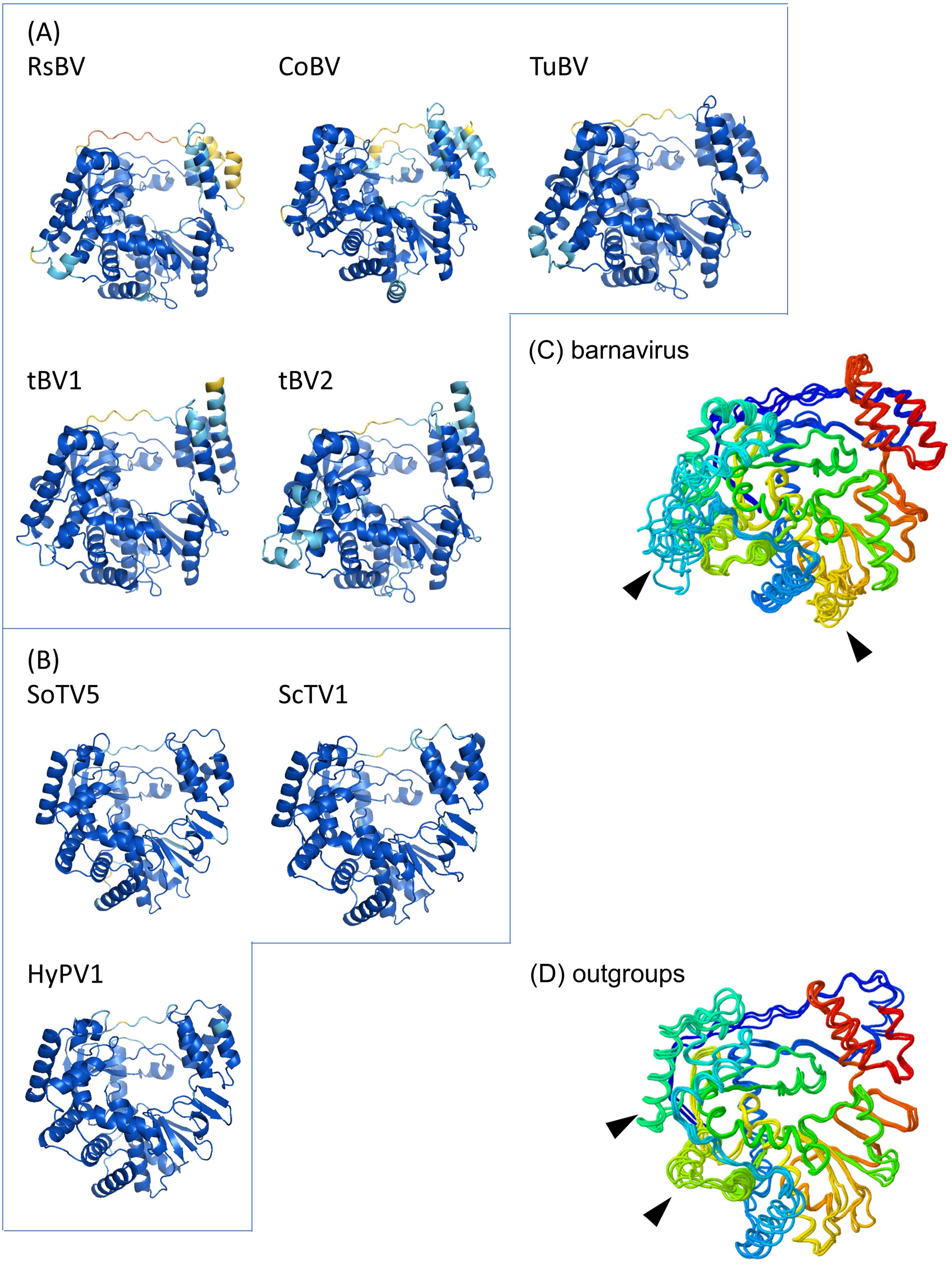
