## Supplementary material for "Integrating structural modeling and divergence dating of RNA-dependent RNA polymerases to resolve the evolutionary history of plant and fungal viruses: from sobemoviruses to sobelivirads": S10 Fig

**S10 Figure** : Structure-guided alignment of RdRp sequences of the 6 barnaviruses (one ICTV-approved and 5 tentative species), 3 outgroups and 4 sobelivirads representative of each other genera. Strictly conserved and similar residues are highlighted (black and grey, respectively), RdRp conserved motifs and homomorphs are outlined (black and grey, respectively). The thumb, index, picky finger domains are outlined pink, magenta and blue violet, respectively.

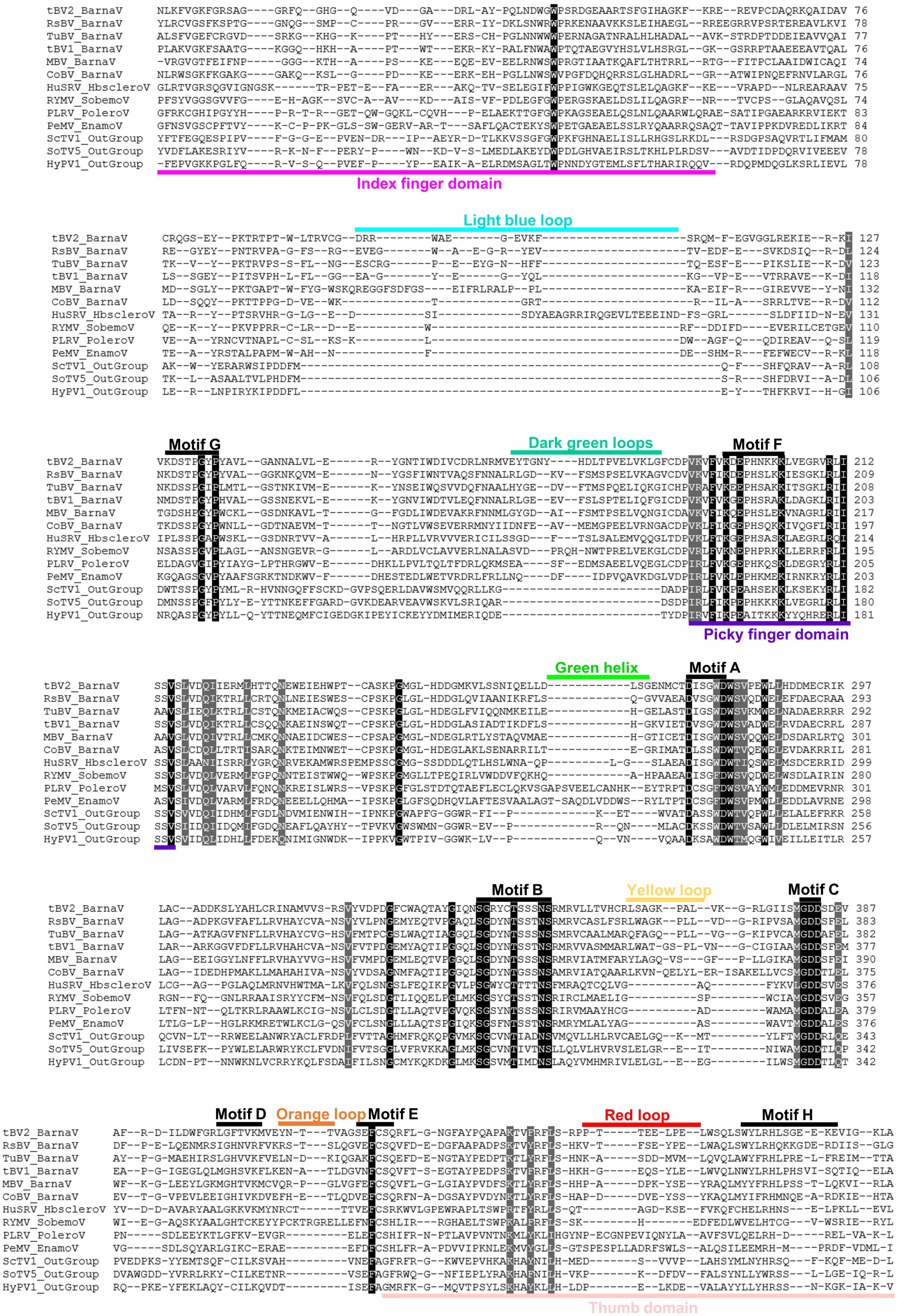
