## Supplementary material for "Integrating structural modeling and divergence dating of RNA-dependent RNA polymerases to resolve the evolutionary history of plant and fungal viruses: from sobemoviruses to sobelivirads": S11 Table

**S11 Table :** Additionnal 99 sequences (genus, name species and accession numbers)

| Genus | Species | Species (old name) | Accession number |
|---|---|---|---|
| Polerovirus | *Polerovirus AEYV* | *African eggplant yellowing virus* | KX856972 |
| | *Polerovirus APVA* | *Allium polerovirus A* | ON565072 |
| | *Polerovirus ARTVB* | *Artemisia virus B* | MT757161 |
| | *Polerovirus BLYV* | *Beet leaf yellowing virus* | LC428352 |
| | *Polerovirus BPV* | *Barleria polerovirus 1* | MW251502 |
| | *Polerovirus BVG* | *Barley virus G* | KT962089 |
| | *Polerovirus BWYV* | *Beet western yellows virus* | NC_004756 |
| | *Polerovirus CBTV1* | *Cotton bunchy top virus 1* | MT966040 |
| | *Polerovirus CBTV2* | *Cotton bunchy top virus 2* | MT966041 |
| | *Polerovirus CDPV* | *Cardamom polerovirus* | BK013145 |
| | *Polerovirus CLDV* | *Cotton leafroll dwarf virus* | NC_014545 |
| | *Polerovirus CNPV* | *Cnidium polerovirus 1* | OP067680 |
| | *Polerovirus CPCSV* | *Chickpea chlorotic stunt virus* | NC_008249 |
| | *Polerovirus CPLRV* | *chickpea leafroll virus* | ON555767 |
| | *Polerovirus CPPV1* | *Cowpea polerovirus 1* | KY364846 |
| | *Polerovirus CPPV2* | *Cowpea polerovirus 2* | KY364847 |
| | *Polerovirus CSPV* | *Cassava polerovirus* | KC505249 |
| | *Polerovirus CTRLV* | *Carrot red leaf virus* | NC_006265 |
| | *Polerovirus CYDVRPV* | *Cereal yellow dwarf virus RPV* | NC_004751 |
| | *Polerovirus DVPV* | *Dregea volubilis polerovirus 1* | MZ965076 |
| | *Polerovirus FBPV* | *Faba bean polerovirus 1* | NC_055495 |
| | *Polerovirus FVPV* | *Foeniculum vulgare polerovirus* | BK059375 |
| | *Polerovirus GRAV* | *Groundnut rosette assistor virus* | MN600000 |
| | *Polerovirus GPOV* | *Grapevine polerovirus 1* | LC507098 |
| | *Polerovirus HEMVA* | *Hemisteptia virus A* | ON416859 |
| | *Polerovirus KMPV* | *Kalanchoe marnieriana polerovirus* | BK059371 |
| | *Polerovirus LABYV* | *Lluffa aphid-borne yellows virus* | KF427701 |
| | *Polerovirus MABYV* | *Melon aphid-borne yellows virus* | NC_010809 |
| | *Polerovirus MAYMV* | *Maize yellow mosaic virus* | MK652150 |
| | *Polerovirus MJVA* | *Mallotus japonicus virus A* | OP122168 |
| | *Polerovirus MYDVRMV* | *Maize yellow dwarf virus RMV* | NC_021484 |
| | *Polerovirus MYFV* | *Miscanthus yellow fleck virus* | MT520166 |
| | *Polerovirus ORMV* | *Ornithogalum virus 5* | MN204612 |
| | *Polerovirus PABYV* | *Pepo aphid-borne yellows virus* | NC_030225 |
| | *Polerovirus PBMYV* | *phasey bean mild yellows virus* | KT962999 |
| | *Polerovirus PDMV* | *Panicum distortion mosaic virus* | LC424839 |
| | *Polerovirus PELRCV* | *pepper leafroll chlorosis virus* | LT220496 |
| | *Polerovirus PEVYV1* | *Pepper vein yellows virus 1* | AB594828 |
| | *Polerovirus PEVYV2* | *Pepper vein yellows virus 2* | HM439608 |
| | *Polerovirus PEVYV3* | *Pepper vein yellows virus 3* | KP326573 |
| | *Polerovirus PEVYV4* | *Pepper vein yellows virus 4* | KU999109 |
| | *Polerovirus PEVYV5* | *Pepper vein yellows virus 5* | KY523072 |
| | *Polerovirus PEVYV6* | *Pepper vein yellows virus 6* | LT559483 |
| | *Polerovirus PEWBVYV* | *pepper whitefly borne vein yellow virus* | MK333461 |
| | *Polerovirus PLAVA* | *Plantago asiatica virus A* | MZ571143 |

| | | | |
|---|---|---|---|
| | *Polerovirus PMPV* | *Piper methysticum polerovirus* | BK059373 |
| | *Polerovirus PNPV* | *Paspalum notatum polerovirus* | BK059372 |
| | *Polerovirus PPEVYV* | *pod pepper vein yellows virus* | MT188667 |
| | *Polerovirus PPOV* | *persimmon polerovirus* | LC488188 |
| | *Polerovirus PTPV* | *Pterostylis polerovirus* | OL471344 |
| | *Polerovirus PUPV* | *Pumpkin polerovirus* | NC_055513 |
| | *Polerovirus SABYV* | *Suakwa aphid-borne yellows virus* | NC_018571 |
| | *Polerovirus SAYV* | *Sauropus yellowing virus* | KJ885302 |
| | *Polerovirus SBCLRV* | *Soybean chlorotic leafroll virus* | OM507197 |
| | *Polerovirus SCYLV* | *Sugarcane yellow leaf virus* | NC_000874 |
| | *Polerovirus SLPV* | *Siratro latent polerovirus* | MK482114 |
| | *Polerovirus SLPSV* | *Sweet potato leaf speckling virus* | - |
| | *Polerovirus SPV* | *Strawberry polerovirus 1* | KM233705 |
| | *Polerovirus STAVB* | *Stellaria aquatica virus B* | OP389993 |
| | *Polerovirus TAPV* | *Trachyspermum ammi polerovirus* | BK059374 |
| | *Polerovirus TCLV* | *Torilis crimson leaf virus* | LT615235 |
| | *Polerovirus TNDV* | *Tobacco necrotic dwarf virus* | - |
| | *Polerovirus TPV1* | *Tobacco polerovirus 1* | MW579552 |
| | *Polerovirus TPV2* | *Tobacco polerovirus 2* | MW579555 |
| | *Polerovirus TRIYSV* | *Triticum yellow stripe virus* | OM829809 |
| | *Polerovirus TVDV* | *Tobacco vein distorting virus* | NC_010732 |
| | *Polerovirus TRIYSV* | *Triticum yellow stripe virus* | OM829809 |
| | *Polerovirus UPOV* | *Ullucus polerovirus 1* | MH978189 |
| | *Polerovirus WCMV* | *White clover mottle virus* | LC192169 |
| | *Polerovirus WLYAV* | *Wheat leaf yellowing-associated virus* | KY605226 |
| | *Polerovirus WYDVGPV* | *Barley yellow dwarf virus - GPV* | FM865413 |
| | *Polerovirus ZABYV* | *zucchini aphid-borne yellows virus* | MK050791 |
| *Enamovirus* | *Enamovirus AGV* | *Ageratum virus 2* | OP660857 |
| | *Enamovirus ALVE* | *Arracacha latent virus E* | MF136435 |
| | *Enamovirus BEV* | *Bean enamovirus 1* | MZ361924 |
| | *Enamovirus BFTEV* | *Birdsfoot trefoil enamovirus 1* | BK010825 |
| | *Enamovirus BPVE* | *Black pepper virus E* | MZ702869 |
| | *Enamovirus CLEV* | *Celmisia lyallii enamovirus* | BK059370 |
| | *Enamovirus GSPEV* | *Ggreen Sichuan pepper enamovirus* | MH323436 |
| | *Enamovirus KSEV* | *Kummerowia striatad enamovirus* | MN814310 |
| | *Enamovirus PEEV* | *Pepper enamovirus* | MG470803 |
| | *Enamovirus PLEV* | *Plantago enamovirus* | MH397359 |
| | *Enamovirus RCEV* | *Red clover enamovirus 1* | MN412742 |
