## Supplementary material for "Integrating structural modeling and divergence dating of RNA-dependent RNA polymerases to resolve the evolutionary history of plant and fungal viruses: from sobemoviruses to sobelivirads": S12 Fig

**S12 Figure** : Phylogenetic tree obtained using Foldtree. Viral species are colored according to the five genera and the outgroups. Two main clades, namely Sobarna- vs. Polena-clades, were revealed clustering sobemo- and barnaviruses vs. poler/mo- and enamoviruses, respectively.

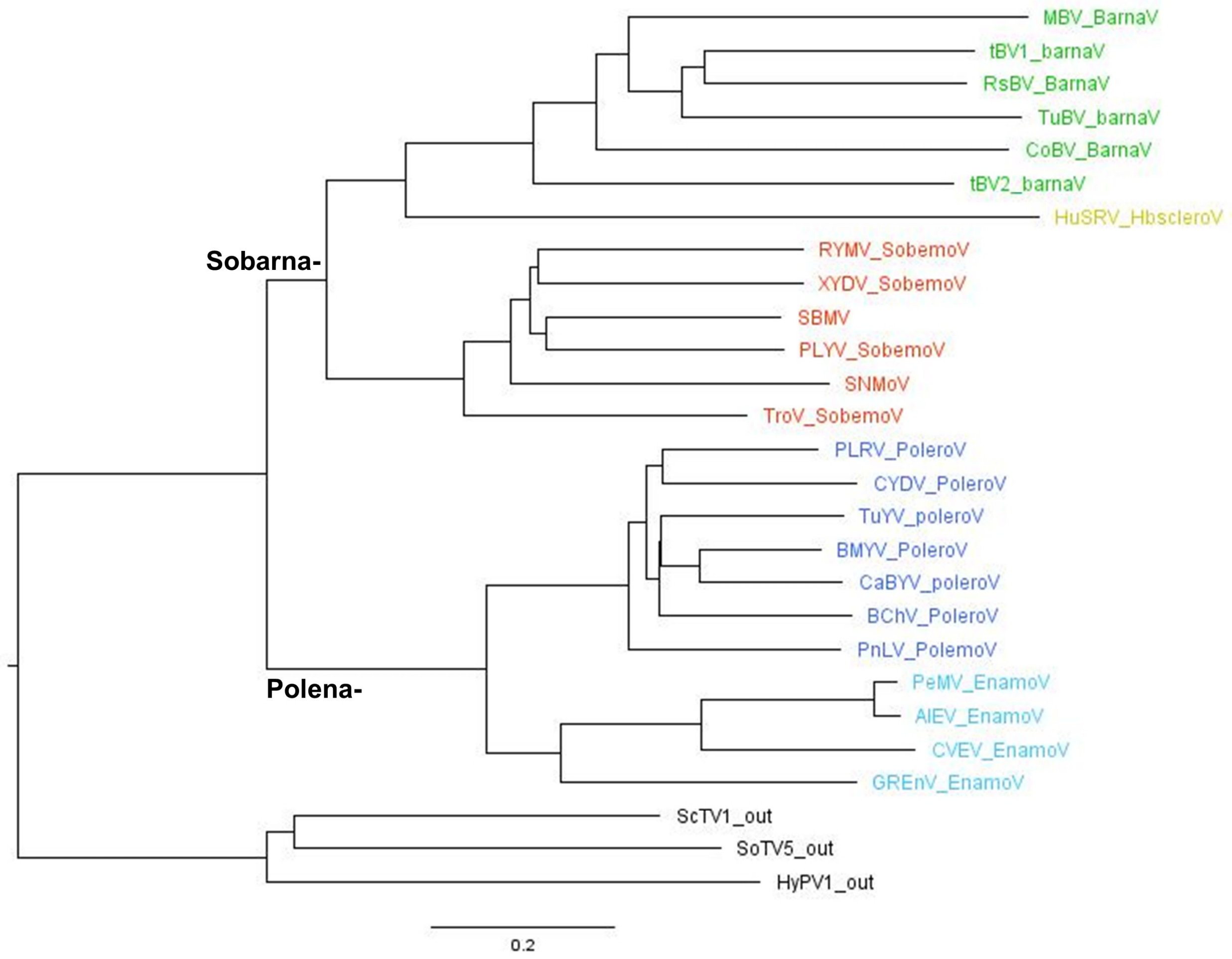
